# Hormone oscillations preserve responsiveness and cross-cell-type coordination of endocrine signaling

**DOI:** 10.64898/2026.08.28.747949

**Authors:** Mark Greenwood, Julia Drube, Carsten Hoffmann, Pulin Li

## Abstract

Living organisms continuously adapt to changing physiological demands by sensing fluctuating concentrations of hormones. This requires hormone signaling pathways to remain responsive over time and discriminate differences in concentration. However, sustained exposure to hormones induces desensitization, which safeguards against overstimulation but may impair cells’ ability to sense future changes in hormone. How the endocrine system circumvents this paradox to preserve responsiveness remains unresolved. Here, using epinephrine-mediated stress signaling as a model, we show that natural ultradian hormone oscillations provide a solution. Oscillatory exposure permits receptor resensitization when hormone levels fall, thereby preserving cellular alertness to future stress and tunability across stimulus intensities. Oscillation also maintains coordination of alertness across diverse cell types by buffering against cell-type variation in receptor kinetics. Ultradian oscillations thus enable endocrine systems to retain protective desensitization while preserving responsiveness and cross-cell-type coordination. Our findings suggest hormone therapies should leverage temporal dynamics to avoid drug-induced tolerance.

## Introduction

Endocrine systems continuously adjust circulating hormone concentrations to signal changing physiological demands, from energy status to reproductive state (Fig. 1A, *left*). Through this stream of communication, target tissues adjust physiology moment to moment, sustaining homeostasis as demands shift. To function effectively, cells must faithfully track these concentration changes over time (‘alertness’) and translate them into physiological responses that scale with demand (‘tunability’) (Fig. 1A, *middle*). Yet, hormone signaling pathways are typically subject to multiple desensitization mechanisms, leading to progressive reduction in responsiveness within minutes of ongoing hormone stimulation (Fig. 1A, *right*)^1–3^. This creates a fundamental conundrum for endocrine signaling: How do cells continue to track and respond to changing hormone concentrations despite desensitization?

**Fig. 1.**
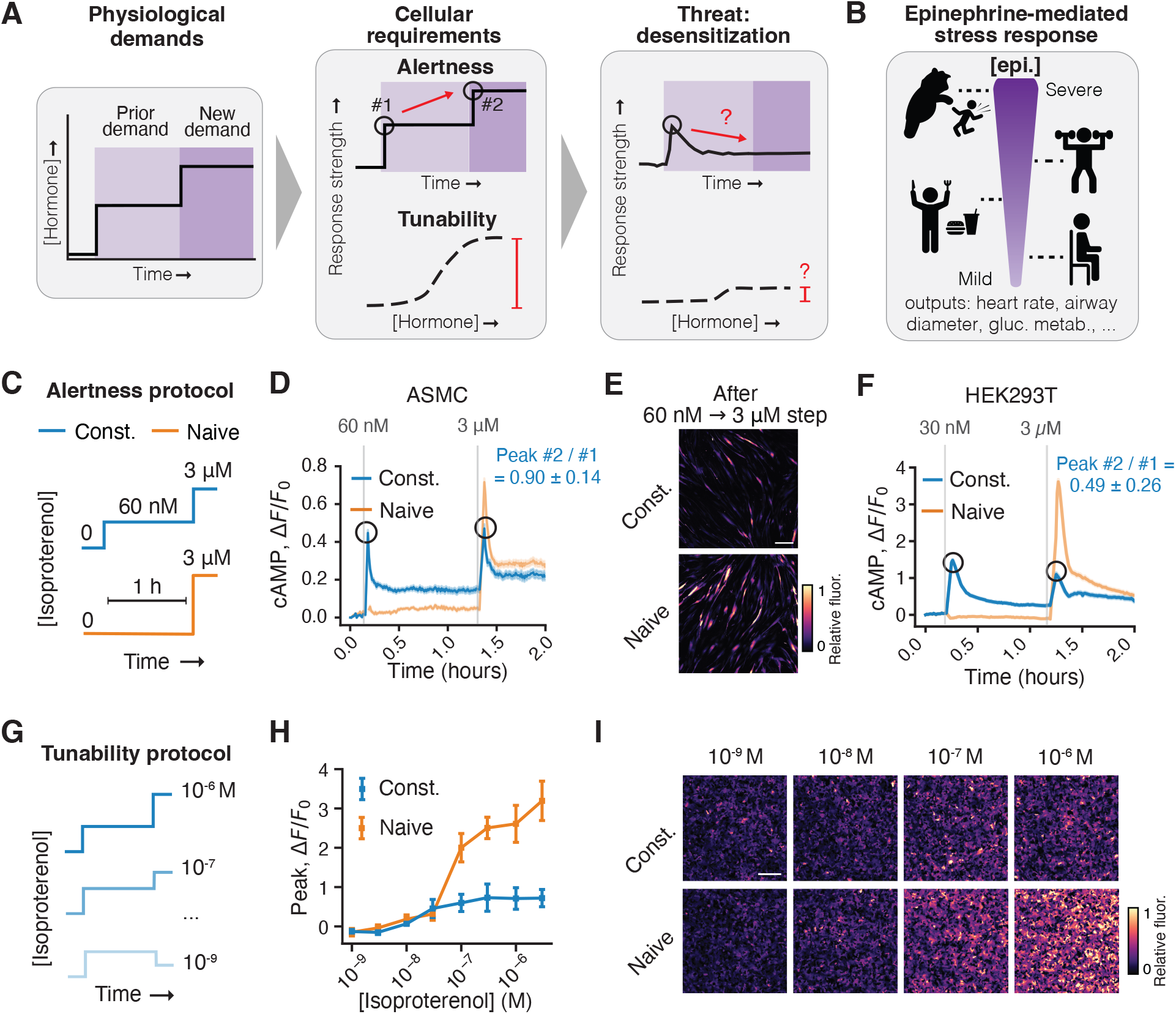
Prior exposure to epinephrine compromises cellular alertness and tunability. (**A**) Physiological demands shift over time (*left*), requiring cells to respond to a new stimulus level from an already-stimulated state (alertness) and grade output across stimulus strength (tunability) (*middle*). Desensitization might compromise both alertness and tunability (*right*). (**B**) Epinephrine regulates homeostasis and stress responses, with levels rising and falling in proportion to physiological demand across a wide range of stress severities. (**C**) Experimental design for assessing alertness. An increase in stress was modeled by transitioning from the approximate EC_50_ (60 nM) to saturating (3 μM) isoproterenol concentration after 1 h (‘constant’ pretreatment condition). A ‘naive’ condition omits the initial low-dose exposure. (**D**) Representative time courses of a cAMP biosensor in airway smooth muscle cells (ASMCs) during the alertness protocol in (C). Signal is reported as Δ*F*/*F₀*, fold change in fluorescence intensity above the baseline. Traces show the mean ± SEM for *n* = 370 (constant) and 428 (naive) cells. The summary statistic shows mean ± SD for *n* = 3 independent experiments. (**E**) Representative biosensor images from (D), showing the peak after the step to 3 µM. Scale bar, 200 μm. (**F**) Representative time courses of the same cAMP biosensor in HEK293T cells during the alertness protocol as in (C), with 30 nM pretreatment. Traces show the mean ± SEM for *n* = 910 (constant) and 511 (naive) cells. The summary statistic shows mean ± SD for *n* = 3 independent experiments. (**G**) Experimental design for assessing tunability. Cells were pretreated constantly at 30 nM, the approximate EC_50_ isoproterenol concentration in HEK293T cells, for 1 h before switching to higher or lower concentrations to construct a dose-response curve. The naive condition steps directly to the high concentration. (**H**) Dose-response curves of peak cAMP (the maximum response after the switch) in HEK293T cells generated by the tunability protocol of (G). Points show mean ± SD for *n* = 3 independent experiments. Paired two-tailed *t* test on the maximal response (*E_max_*) between constant and naive conditions: *P* = 0.0077. (**I**) Representative images from (H), showing the peak after the step change in concentration. Scale bar, 200 μm.

Despite its apparent drawbacks, desensitization protects tissues from pathological overstimulation. Epinephrine signaling, a central regulator of mammalian homeostasis and stress responses, is a classic example. Excessive epinephrine signaling contributes to cardiac hypertrophy^4–8^, and β-arrestins, which mediate receptor desensitization, simultaneously initiate alternative signaling pathways that counteract overstimulation toxicity^9^. At the same time, epinephrine is continuously present in circulation, and its concentration scales dynamically with the intensity of physiological stress to drive responses across tissues, including changes in heart rate, glucose metabolism, and airway tone (Fig. 1B)^10–12^. Endocrine control must therefore reconcile two competing requirements: retaining the protective effects of desensitization while preserving the capacity to respond to subsequent changes in hormone concentration.

Here, we show that physiological ultradian hormone oscillations reconcile desensitization with sustained responsiveness, and in doing so keep responsiveness coordinated across cell types. Combining quantitative live-cell imaging, genetic perturbations, and mathematical modeling, we find that in epinephrine signaling, receptor desensitization erodes alertness and tunability, whereas oscillations preserve them by creating recurring low-hormone intervals that permit receptor resensitization. Compared with alternative strategies, oscillation preserves alertness and tunability more consistently across hormone concentrations, and improves coordination between different cell types with divergent receptor kinetics. These findings establish hormone oscillations as a design principle by which endocrine signaling reconciles receptor desensitization with sustained and coordinated responsiveness, and suggest restoring oscillations as a strategy for hormone therapies.

### Prior exposure to epinephrine compromises cellular alertness and tunability

Faithful tracking of dynamic hormone signals requires alertness and tunability, so that mild and severe stimuli produce proportionally different outputs. We first asked whether cells can maintain alertness and tunability after prior stimulation, since intensifying stress raises circulating epinephrine above its prior level rather than from zero. We exposed cells to a low concentration of the epinephrine analog isoproterenol before raising it to a saturating concentration without washout (‘constant pretreatment’ condition; Fig. 1C). In parallel, cells without any prior exposure directly received the saturating concentration (‘naive’ condition). The naive condition therefore defines the maximal response of an unadapted cell, whereas any reduction in the constant pretreatment condition reflects a loss of alertness.

We focused on airway smooth muscle cells (ASMCs), which utilize tonic epinephrine signaling to maintain homeostatic airway tone and acute epinephrine signaling to dilate airways during stress^13^. In ASMCs, epinephrine signals through β-adrenergic receptors to stimulate cAMP production^14^, and we confirmed that ADRB2 is the predominant subtype expressed endogenously (table S1). To monitor signaling, we expressed a genetically encoded biosensor for cAMP^15^, which exhibited a dose-dependent response to isoproterenol (Fig. S1).

Both low and saturating isoproterenol concentrations elicited the expected transient increase in cAMP, peaking within minutes before attenuating, consistent with desensitization (Fig. 1D). We quantified alertness as the ratio of the peak cAMP response to the high concentration relative to the peak response to the preceding low concentration. Because the initial concentration (60 nM) approximates the EC_50_ and the second (3 µM) is saturating, a fully alert cell is expected to produce a ratio close to 2, whereas ratios at or below 1 indicate a failure to respond to the increase in stress.

Increasing isoproterenol from 60 nM to 3 µM produced essentially no additional signaling (Fig. 1D), whereas naive cells exposed directly to 3 µM generated a much larger response (Fig. 1D and E; Fig. S2A and B; movie S1). Thus, cells adapted to a constant low level of stress fail to accurately sense subsequently elevated stress. Furthermore, we observed loss of alertness in HEK293T cells (Fig. 1F; Fig. S2C and D; movie S2), which also predominantly express the β2 receptor subtype (table S1), indicating that the loss of alertness is not cell-type specific.

We next asked whether cells pre-exposed to epinephrine also lose the ability to quantitatively adjust their responses according to ligand concentrations (tunability). To test this, we measured responses across a wider range of isoproterenol concentrations following pretreatment. HEK293T cells were pretreated at a fixed concentration and then shifted to a series of higher or lower concentrations (Fig. 1G). Compared to naive cells, pretreated cells exhibited a markedly compressed dose-response curve, indicating a substantial loss of tunability (Fig. 1H and I). Together, these results demonstrate that prior exposure of cells compromises alertness and tunability in response to elevated stress potentially due to ligand-induced signaling desensitization.

### Receptor desensitization causes loss of alertness and tunability

Signaling desensitization can occur through receptor inactivation or internalization^16–21^, reduced second-messenger synthesis or accelerated degradation^22–25^, and transcriptional remodeling^26,27^. Although each of these can contribute to desensitization, we asked which mechanism functionally drives the loss of alertness and tunability (Fig. 2A). Bypassing the receptor with forskolin, which directly activates the cAMP-producing enzyme adenylyl cyclase, fully prevented the loss of alertness, and by extension tunability (Fig. 2B and C; Fig. S3A and B), placing the functional defect at the receptor. Consistent with a receptor-level origin, surface ADRB2 labeled with membrane impermeable HaloTag ligand fell upon ligand stimulation and partially recovered upon washout, indicating desensitization and resensitization occur at the receptor (Fig. 2D and Fig. S3C).

**Fig. 2.**
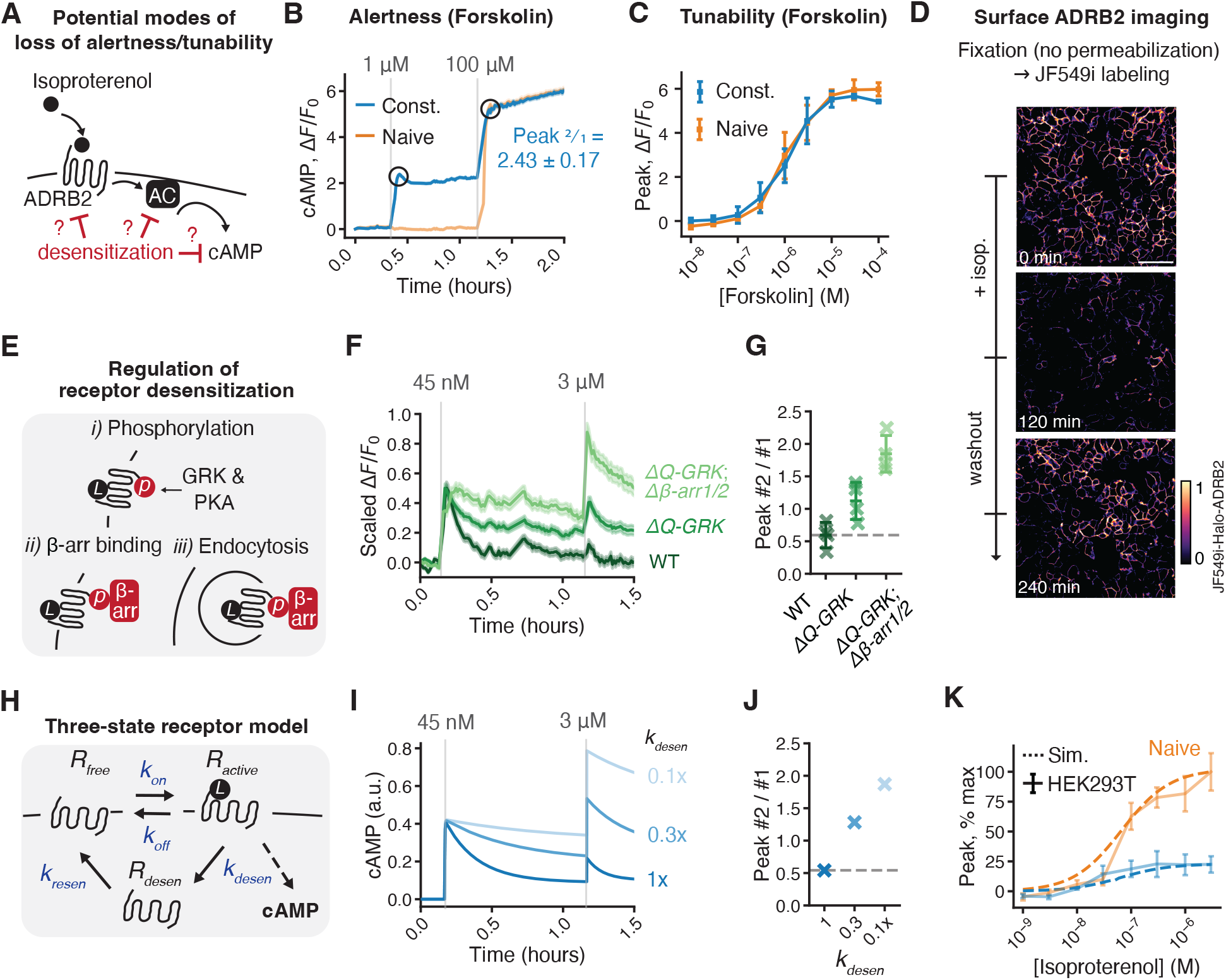
Receptor desensitization and resensitization kinetics capture loss of alertness and tunability. (**A**) Isoproterenol binding to ADRB2 activates adenylyl cyclase (AC) to drive cAMP production. Loss of alertness could arise at multiple levels: ADRB2 desensitization, feedback inhibition of AC, or accelerated degradation of cAMP. (**B**) Representative cAMP time course during the alertness protocol in HEK293T cells stimulated with forskolin, under constant pretreatment or naive condition. Traces show the mean ± SEM for *n* = 737 (constant) and 262 (naive) cells. The summary statistic shows mean ± SD for *n* = 3 independent experiments. (**C**) Dose-response curves of peak cAMP in HEK293T cells generated by the tunability protocol using forskolin. Forskolin bypasses the receptor to activate cAMP synthesis directly via adenylyl cyclase (AC). Points show mean ± SD for *n* = 3 independent experiments. Paired two-tailed *t* test on the maximal response (*E_max_*) between constant and naive conditions: *P* = 0.22. (**D**) Representative images of JF549i-Halo-ADRB2 in HEK293T cells at baseline (naive), after isoproterenol treatment (120 min), and following washout (240 min). Scale bar, 100 μm. (**E**) Receptor regulation cascade: ligand bound receptor is phosphorylated by GRK and PKA (*i*), recruits β-arrestin (*ii*), and is internalized (*iii*). (**F**) Representative cAMP time courses in WT HEK293, *ΔQ-GRK*, and *ΔQ-GRK* / *Δβ-arr1/2* cells under the alertness protocol. Traces show the mean ± SEM for *n* = 101, 154, and 123 cells (WT, ΔQ-*GRK*, *ΔQ-GRK* / *Δβ-arr1/2*, respectively) from one representative experiment. (**G**) The ratio of peak heights (peak #2 / #1) for the genotypes in (F). Crosses, individual experiments; bars mean ± SD; *n* = 4 independent experiments. One-way ANOVA (*P* < 0.001) with Tukey’s HSD post-hoc test: WT vs Δ*Q-GRK*, *P* = 0.043; WT vs *ΔQ-GRK* / *Δβ-arr1/2*, *P* < 0.001; Δ*Q-GRK* vs *ΔQ-GRK* / *Δβ-arr1/2*, *P* = 0.008. (**H**) A three-state receptor model describes β-adrenergic receptor kinetics, including ligand binding (*R_free_* ↔ *R_active_*), receptor desensitization (*R_active_* → *R_desen_*) and resensitization (*R_desen_* → *R_free_*), with cAMP production downstream of the active receptor state (*R_active_*). (**I**) Simulated cAMP time courses demonstrating the effect of varying *k_desen_* during the alertness protocol, which recapitulates (F). (**J**) The ratio of peak heights (peak #2 / #1) for different *k_desen_* rates simulated in (I), which recapitulates (G). (**K**) Comparison of experimentally measured tunability in HEK293T cells with the model simulation, under naive and constant 30 nM pretreatment conditions. Points show mean ± SD for *n* = 3 independent experiments.

We next asked which regulatory machinery drives the loss, and whether disrupting it restores alertness. We genetically slowed ADRB2 desensitization by disrupting two sequential regulatory steps, GRK-mediated phosphorylation and β-arrestin binding^20,21^ (Fig. 2E). We used HEK293 cells lacking GRK2, 3, 5, and 6 (Δ*Q-GRK*)^28^, and cells additionally lacking both β-arrestin isoforms (Δ*Q-GRK* / Δ*β-arr1/2*)^29^. Both perturbations slowed cAMP attenuation after isoproterenol stimulation, confirming reduced desensitization, with Δ*Q-GRK* / Δ*β-arr1/2* slowing it further than Δ*Q-GRK* alone, while leaving the isoproterenol EC_50_ largely unchanged (Fig. S4). Consistent with this graded loss of desensitization, alertness recovered modestly in Δ*Q-GRK* cells and substantially in Δ*Q-GRK* / Δ*β-arr1/2* cells (Fig. 2F and G). Therefore, the rate of receptor desensitization, through GRKs and β-arrestin, sets the alertness and tunability of cells.

To ask if receptor desensitization and resensitization are sufficient to account for alertness and tunability, we built a minimal model of receptor signaling. Receptors interconvert between an unbound (*R_free_*) and a ligand-bound, cAMP-producing active state (*R_active_*), with active receptors entering a signaling-incompetent desensitized state (*R_desen_*) before resensitizing to the free state (Fig. 2H). Fit to ASMC and HEK293 cAMP dynamics measured during the alertness protocols (Fig. 1D and F), and with no further fitting thereafter, the model reproduced alertness across desensitization rates (Fig. 2I and J; Fig. S5; tables S2 and S3) and the loss of tunability (Fig. 2K). Together, these results establish receptor desensitization and resensitization as the determinants of cellular alertness and tunability.

### Oscillation preserves alertness and tunability

A critical difference between physiological and experimental stimulation may provide the solution to the alertness and tunability problem. Most hormones *in vivo* exhibit natural ultradian oscillations^30,31^; epinephrine oscillates with periods of approximately 0.5–2 hours, and the amplitude of these oscillations increases with physiological stress^32–34^. This oscillation creates a transient period of low ligand, which could potentially allow cells to reset their available receptor pool. Therefore, we hypothesized that these oscillations could preserve cellular alertness and tunability, and allow cells to better adapt to physiological demands.

To test this idea, we first applied our model to compare responses following either constant or oscillatory pretreatment, in which the ligand alternated between zero and the same concentration, before increasing to a higher concentration (Fig. 3A). The model predicted that oscillatory pretreatment preserves alertness: the ratio of cAMP peak responses from the final to the initial concentration was 1.67 under oscillations and 0.99 under constant pretreatment (Fig. 3B). This advantage persisted across a broad range of oscillation parameters, with lower peak, lower trough concentrations, and shorter duty cycles favoring improvement in alertness (Fig. S6), and persisted when cAMP metabolism and receptor turnover were incorporated into the model (Fig. S7; table S4).

**Fig. 3.**
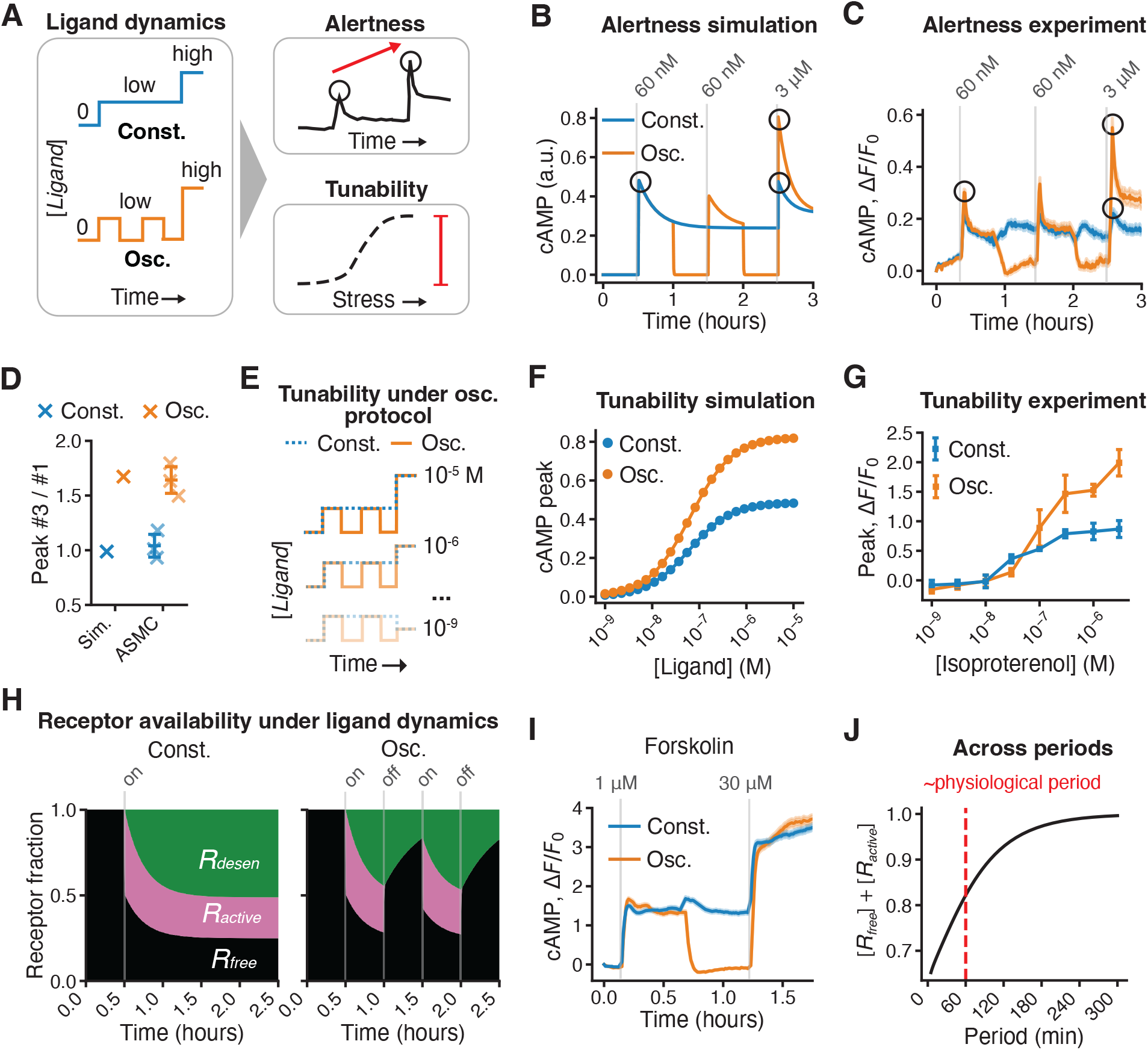
Epinephrine oscillation preserves the alertness and tunability of the stress response. (**A**) Assessing how exposure to either constant or oscillatory low-dose pretreatment affects alertness and tunability to subsequent changes in hormone concentration. (**B**) Simulated cAMP time courses for the alertness protocol under constant or oscillatory pretreatment. (**C**) Representative cAMP time courses during the alertness protocol in ASMCs, under constant or oscillatory pretreatment. Traces show the mean ± SEM for *n* = 502 (constant) and 454 (oscillatory) cells. (**D**) The ratio of peak heights (peak #3 / #1) for the simulation in (B) and experimental measurements in (C) under constant and oscillatory pretreatment. Crosses, individual experiments; bars, mean ± SD; *n* = 3 independent experiments. Constant vs. oscillation (experiment): paired two-tailed *t* test, *P* = 0.001. (**E**) Comparing tunability under constant vs. oscillatory conditions. Cells were pretreated with constant or oscillatory isoproterenol at 30 nM before stepping to a range of ligand concentrations. (**F**) Simulated dose-response curves of peak cAMP under the tunability protocol in (E), with constant or oscillatory pretreatment. (**G**) Dose-response curves of peak cAMP in HEK293T cells under the tunability protocol in (E), with oscillatory or constant pretreatment. Points show mean ± SD for *n* = 3 independent experiments. Paired two-tailed *t* test on the maximal response (*E_max_*) between constant and oscillatory pretreatment: *P* = 0.026. (**H**) Receptor state distribution during pretreatment in simulation. (**I**) Representative cAMP time courses during the alertness protocol in HEK293T cells with forskolin, under constant or oscillatory pretreatment. Traces show the mean ± SEM for *n* = 509 (constant) and 466 (oscillatory) cells. (**J**) The simulated available receptor (*R_free_* + *R_active_*) at the end of an oscillation cycle as a function of period length. The red line marks the approximate period length of physiologically observed oscillation.

We next tested the model prediction experimentally. Consistent with the model, oscillatory pretreatment preserved cellular alertness to subsequent increases in agonist concentration (Fig. 3C). In ASMCs, the ratio of cAMP peak responses to the final versus initial concentration increased from 1.04 ± 0.10 under constant isoproterenol pretreatment to 1.64 ± 0.12 under oscillations (Fig. 3D). HEK293T cells exhibited a qualitatively similar improvement of alertness under oscillations (Fig. S8A and B).

Next, we asked whether oscillation also preserves tunability. We simulated a change in stress by abruptly increasing or decreasing the ligand concentration across a range of values (Fig. 3E), and found that oscillatory pretreatment substantially increased the maximal efficacy of the dose-response curve, thereby expanding the tunability of the response (Fig. 3F). We tested this prediction experimentally in HEK293T cells by measuring responses to concentration changes spanning a wide range of isoproterenol concentrations. Consistent with the simulations, oscillatory pretreatment expanded the dose-response curve, restoring tunability compared with constant pretreatment (Fig. 3G). Together, these results demonstrate that physiological oscillations preserve both alertness to changing stress levels and tunability across stimulus strengths, enabling cells to maintain responsiveness despite ongoing receptor desensitization.

### Oscillation preserves alertness and tunability by restoring receptor availability

How does oscillation preserve alertness and tunability? Examination of the receptor species under oscillation simulations revealed that oscillation allowed free receptors to recover during each ligand-off phase, periodically replenishing the pool of available receptors (Fig. 3H, *right*). In contrast, constant stimulation drove receptors into an equilibrium with a smaller pool of free receptors (Fig. 3H, *left*). This leaves more free receptors available under oscillation to respond to changes in stress. Consistent with this receptor-level mechanism, bypassing the receptor with forskolin abolished the benefits provided by oscillation (Fig. 3I; Fig. S8C and D).

The pool-refilling mechanism predicts that oscillation should help only once the ligand-off phase is long enough for receptors to recover. Varying the oscillation period in the model, we found that the available, signaling-competent receptor pool (*R_free_* + *R_active_*) stayed depleted at short periods, then recovered and saturated near full availability beyond approximately 180 min (Fig. 3J). Notably, the physiological period of approximately 60 min^32–34^ falls on the shoulder of the curve, where the ligand off-phase is comparable to 1 / *k_resen_* for ASMCs (30.3 min), so oscillation restores approximately 80 % of the receptor pool without requiring the longer periods needed for complete recovery. Oscillation therefore preserves alertness and tunability by periodically refilling the available receptor pool, and the physiological period is long enough to facilitate most of the recovery.

### Oscillation preserves responsiveness more consistently across stress levels than alternative strategies

If oscillation preserves responsiveness by restoring available receptors, could cells achieve the same outcome through other means without oscillation? In principle, any condition that maintains a large pool of available receptors during constant stimulation could preserve alertness and tunability, whether by lowering the operating ligand concentration, increasing the receptor abundance, or combining both strategies. We used our model to compare these three alternative strategies with oscillation (Fig. 4A).

**Fig. 4.**
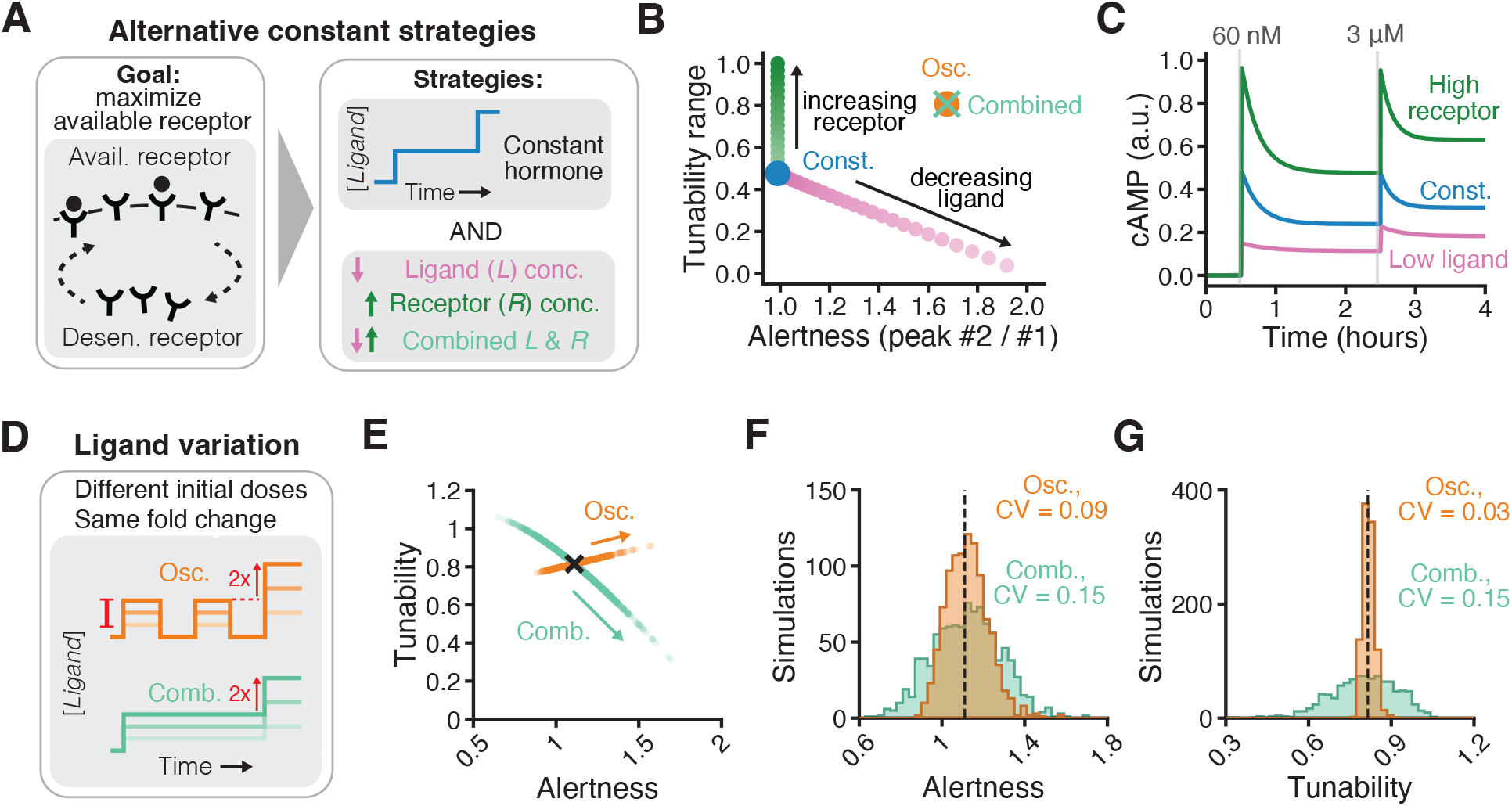
Oscillation preserves responsiveness consistently across stress levels. (**A**) Comparison between oscillation and alternative strategies that operate under constant pretreatment. Alternative strategies, including lower operating ligand (*L*) concentrations, increased receptor (*R*) abundance, or both combined, are designed to maximize available surface receptors after pretreatment. (**B**) Changes in alertness and tunability in response to gradually decreasing ligand concentrations (*pink*) or increasing receptor levels (*green*) under constant stimulation, benchmarked against constant (*blue*) and oscillatory (*orange*) conditions without parameter tuning. Only by simultaneously decreasing ligand concentration and increasing receptor level (*cross*) could the constant condition be tuned to match the performance achieved by oscillations. (**C**) Simulated cAMP time courses during the alertness protocol under the low-ligand only or high-receptor only strategies, benchmarked against constant pretreatment without fine tuning. (**D**) Assessing how variation in operating ligand concentration affects alertness and tunability under the oscillatory or combined strategy. The initial concentration varies between cells, while the fold change of the step is held constant. (**E**) Alertness and tunability across a scan of operating dose under the oscillatory or combined strategy. *n* = 1,000 simulated cells. Arrows point to decreasing operating dose. Cross, the matched reference condition. (**F** and **G**) Distributions of alertness (F) and tunability (G) for the simulations in (E). Dashed lines, values at the reference operating dose. CV, coefficient of variation.

Lowering the operating ligand concentration while keeping the same fold change improved alertness by reducing receptor desensitization, but proportionally diminished tunability (Fig. 4B and C). Lowering ligand concentration confines cells to the lower portion of the dose-response curve, thereby compressing the range over which responses could vary. In contrast, increasing receptor concentration improved tunability by expanding the absolute dose-response range, but left alertness unchanged (Fig. 4B and C). This is because signal responses to the first and second stimuli increased proportionally, leaving the relative difference between the peaks (*i.e.*, alertness) unchanged. Combining a lower ligand concentration with greater receptor abundance could match the performance achieved by oscillation (Fig. 4B, *cross*). However, when the operating ligand concentration varied (Fig. 4D), the combined strategy produced greater variation in alertness and tunability than did oscillation (Fig. 4E to G). Thus, as hormone concentrations vary with physiological demand, oscillation preserves responsiveness more consistently than alternative strategies.

### Oscillation coordinates responsiveness across cell types

The physiological functions of systemic hormones depend not only on the responsiveness of individual cell types but also on the coordination of responsiveness across diverse tissues. During acute stress, for example, epinephrine regulates a wide range of physiological processes, including cardiac contractility, vascular tone, and airway tone^10,12,13^. Proportional changes in these outputs among distinct cell types ensure oxygen delivery to peripheral tissues during the fight-or-flight response^35^. Yet cell types need not desensitize and resensitize at the same rates^36–38^. Consequently, cell types exposed to the same hormone concentration could lose responsiveness unequally, and their outputs would diverge. We therefore asked whether hormone oscillations could preserve coordination of responsiveness across cell types.

To establish if, and by how much, cell types differ in their kinetics, we compiled experimentally measured desensitization and resensitization rates from diverse cell types. These measurements span immortalized cell lines and primary cells across multiple organs and species, from chick ventricular myocytes to rodent cortical neurons (tables S5 and S6). In parallel, to observe the consequence of differences in rates, we computationally mapped the amount of available receptor under constant pretreatment across ranges of desensitization and resensitization rates (Fig. 5A). This map showed simulated cells occupying different positions within the reported range retain markedly different fractions of available receptors after constant pretreatment (Fig. 5A). Literature rates of different cell types varied over approximately 2 orders of magnitude, occupying a region of the map within which available receptor ranges from 0.4 % to 96.7 % of total. These results suggest that due to substantial differences in the receptor kinetics, different cell types would have drastically different levels of alertness under constant hormone exposure, and consequently would lose the coordination in the relative strength of the responses to the same hormone input.

**Fig. 5.**
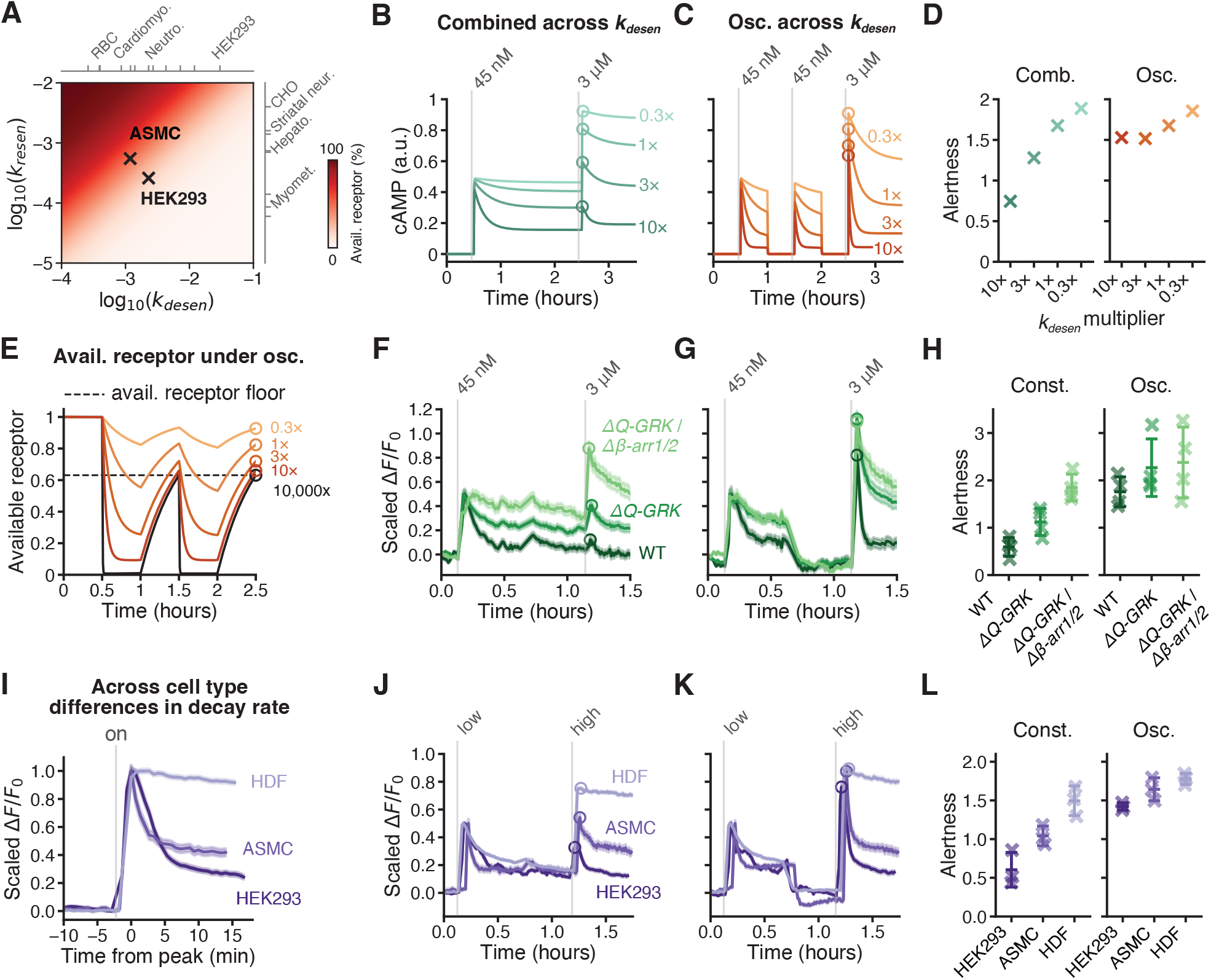
Oscillation coordinates responsiveness across cell types. (**A**) Simulated available receptor (*R_free_* + *R_active_*) under constant pretreatment, across a scan of desensitization (*k_desen_*) and resensitization (*k_resen_*_)_ rates (sec⁻¹). Crosses mark the fitted parameters for ASMCs and HEK293 cells; ticks mark literature-reported desensitization and resensitization rates, respectively, across cell types, with representative cell types labeled. (**B** and **C**) Simulated cAMP time courses during the alertness protocol under the combined constant strategy (B) or oscillatory pretreatment (C), with *k_desen_* varied from its fitted ASMC value. Circles, peak response to the final concentration. (**D**) Alertness (final peak / first peak) for the simulations in (B) and (C). (**E**) Simulated available receptor (*R_free_* + *R_active_*) during oscillatory pretreatment for the *k_desen_* rates in (C) and at maximal desensitization (*10,000x*). Circles, available receptor immediately before the step to the final concentration. Dashed line, the floor in available receptor set by resensitization during the ligand-off phase, independent of *k_desen_*. (**F** and **G**) Representative cAMP time courses in WT HEK293, Δ*Q-GRK*, and Δ*Q-GRK* / Δ*β-arr1/2* cells during the alertness protocol under constant (F) or oscillatory (G) pretreatment. Signal is reported as scaled Δ*F*/*F₀*, normalized to the first peak. Traces show the mean ± SEM for *n* = 101, 154, and 123 cells (WT, Δ*Q-GRK*, Δ*Q-GRK* / Δ*β-arr1/2*, respectively) in (F) and *n* = 81, 131, and 126 cells in (G), from one representative experiment. Circles, peak response to the final concentration. (**H**) Alertness (peak #2 / #1) for the genotypes in (F) and (G). Crosses, individual experiments; bars, mean ± SD; *n* = 4 independent experiments. One-way ANOVA: constant, *P* < 0.001; oscillatory, *P* = 0.32. (**I**) Scaled cAMP time courses following the response to isoproterenol in HEK293 cells, ASMCs, HDFs, aligned to the peak (*t* = 0), showing cell-type differences in apparent cAMP decay rate. Traces show the mean ± SEM for *n* = 774, 393, and 656 cells (HEK293 cells, ASMCs, HDFs, respectively). (**J** and **K**) As in (F and G), for HEK293T cells, ASMCs, and HDFs. Traces show the mean ± SEM for *n* = 611, 301, and 766 cells (HEK293T cells, ASMCs, HDFs, respectively) in (J) and *n* = 774, 393, and 656 cells in (K). (**L**) As in (H), for the cell types in (J) and (K). *n* = 3 independent experiments for each cell type. One-way ANOVA: constant, *P* = 0.003; oscillatory, *P* = 0.013.

To test whether oscillation can help coordinate responsiveness among different cell types, we modeled distinct cell types by independently varying receptor desensitization and resensitization rates in simulations. Under the combined (low ligand and high receptor) constant strategy, cells with different desensitization rates had similar magnitudes of peak responses during pretreatment, but diverged drastically when the hormone concentration increased (Fig. 5B), so that cells had very different degrees of alertness (Fig. 5D and Fig. S9). In contrast, under oscillatory pretreatment, cells arrived at similar peak responses to the acute stress (Fig. 5C) and had more similar alertness (Fig. 5D and Fig. S9). This result follows from the mechanism: during oscillation, each fall in ligand allows desensitized receptors to recover, so that available receptor returns to above a floor that is set by both the resensitization rate and the duration of the low-ligand phase, independent of how rapidly receptors desensitize (Fig. 5E). Even cells that desensitize maximally during the high-ligand phase re-enter the following cycle at this floor, thereby buffering differences in *k_desen_*. Consistent with recovery during the low-ligand phase being rate-limiting, the buffering effect held across cell types with faster resensitization rates, though not slower ones (Fig. S9).

To test this prediction, we utilized the Δ*Q-GRK* and Δ*Q-GRK* / Δ*β-arr1/2* HEK293 cells, which desensitize slower than WT cells. Under constant pretreatment, the three genotypes reached different degrees of alertness (Fig. 5F and H). Under oscillatory pretreatment, their alertness was indistinguishable (Fig. 5G and H). Oscillation therefore coordinates responsiveness between cells that differ in desensitization kinetics.

Finally, we asked whether oscillation also buffers the naturally occurring differences between cell types. To span a wide range of kinetics, we included human dermal fibroblasts (HDFs), as we found they desensitize unusually slowly (Fig. 5I). Under constant pretreatment, these cell types reached different degrees of alertness (Fig. 5J and L). Oscillatory pretreatment reduced the spread between them, so that the cell types had similar alertness (Fig. 5K and L). Thus, a single oscillating input can elicit proportionate responses from heterogeneous tissues without requiring that their receptors behave alike.

## Discussion

The endocrine system faces a fundamental conundrum: sustained hormone signaling drives cellular desensitization, yet cells must remain responsive to changing demand over time. Here, using the epinephrine-mediated stress response as an example, we show that natural ultradian oscillations of hormone concentration resolve this tension, preserving cell’s alertness and tunability to changes in stress. Further, oscillation facilitates coordination among diverse cell types, such that the physiological outputs from different organs remain proportional to the stress.

Our finding introduces a novel functional role for oscillatory signaling dynamics. Prior work has largely emphasized how oscillations are decoded into different outputs, expanding the repertoire of responses a single pathway can produce, as reported for intracellular p53^39^, ERK^40^, and NF-κB^41,42^, and for extracellular signals including dopamine^43^, cortisol^44^, and GnRH^45^. Our results reveal a distinct function: rather than shaping the response to the current signal, oscillation preserves a cell’s capacity to respond to future changes in signal intensity, maintaining alertness and tunability to physiological demands not yet encountered.

The preserved responsiveness we observed under oscillatory epinephrine signaling could represent a principle that extends beyond β-adrenergic receptors. Ultradian hormone oscillations are the rule rather than the exception, from insulin to the sex hormones^30,31^, and our mechanism requires only that receptors desensitize and resensitize on timescales matched to the oscillation. Many GPCRs share this property, as might some receptor tyrosine kinases such as the insulin receptor^46,47^. Whether an analogous principle operates for receptor classes that regulate sensitivity by other means, such as nuclear hormone receptors that lack membrane desensitization, remains open. Testing across these receptor classes and diverse cell types will define the scope of the effect and may help explain why hormone oscillation is so pervasive.

Our findings identify temporal structure as a mechanism to coordinate responses across cell types. Within tissues, cells commonly coordinate through intercellular communication, including gap junctional, juxtracrine, or paracrine signaling, all of which depend on proximity. Alternatively, cells can coordinate responses through shared intracellular architectures, such as fold-change detection motifs, which buffer differences in signaling properties among cells of a given type^48–50^. We found that oscillations offer a distinct strategy, coordinating responses even across different cell types. Because coordination is encoded in the hormone signal itself, it requires neither proximity between cells nor specialized network architecture within them. Hormone oscillations may therefore be a general strategy for maintaining coherent responses among cells with distinct signaling properties, irrespective of their location within a tissue or across the body.

β-adrenergic agonists are among the most widely used bronchodilators, yet their sustained use can diminish the responsiveness they are intended to provide^51,52^. Although our experiments operate on shorter timescales than clinical usage, they capture receptor desensitization in the same receptor system and the ASMCs in which tolerance develops^53,54^. Our findings therefore raise the possibility that drug delivery could be optimized not only in dose but also in time: delivery regimens that incorporate periods of low agonist exposure may allow receptor resensitization and preserve subsequent responsiveness. More broadly, engineering signals in time may provide a way to control not only how cells respond now, but how responsive they remain to what comes next.

## Acknowledgments

We thank Gavin Schlissel for critical feedback on the manuscript and Michael Elowitz for helpful discussion on the project.

## Author contributions

Conceptualization: MG & PL

Methodology: MG & PL

Investigation: MG

Visualization: MG

Funding acquisition: MG & PL

Project administration: MG & PL

Supervision: PL

Writing – original draft: MG

Writing – review & editing: MG & PL

Writing – review: JD & CH

JD & CH generated the Δ*Q-GRK* and Δ*Q-GRK* / Δ*β-arr1/2* HEK293 cell lines.

## Data, code, and materials availability

Data and model code will be deposited in public repositories with a citable DOI provided upon revision. Plasmids generated in this study are available from the corresponding author.

## Supplementary Materials

Materials and Methods

Figs. S1 to S9

Tables S1 to S6

Movies S1 and S2

## Materials and Methods

### Cells and cell culture

ASMCs (LifeLine Cell Technology, FC-0059) were cultured in VascuLife basal medium (LifeLine Cell Technology, LM-0002) supplemented with SMC LifeFactors Kit (LifeLine Cell Technology, LM-1040) and 1 % penicillin-streptomycin (Gibco, 15140122). Adult HDFs (ATCC, PCS-201-012) were cultured in Fibroblast Basal Medium (ATCC, PCS-201-030) supplemented with Fibroblast Growth Kit-Low Serum (ATCC, PCS-201-041) and 1 % penicillin-streptomycin. ASMCs and HDFs were passaged once after being received from the manufacturers before use in experiments. The Δ*Q-GRK*, and Δ*Q-GRK* with *β-arrestin 1/2* (Δ*Q-GRK* / *Δβ-arr1/2*) knockout mutants were made previously and are in a HEK293 background ^28,29^. HEK293T (RRID:CVCL_0063) and HEK293 (RRID:CVCL_0045) cells were cultured in FluoroBrite DMEM (Gibco, A1896701) supplemented with 10 % fetal bovine serum (Takara Bio, 631367) and 1 % penicillin-streptomycin-glutamine (Gibco, 10378016). For all cell types and experiments, we cultured cells to confluence in 24- or 96-well clear bottom imaging plates (Cellvis, P24-1.5P and P96-1.5P) treated with 50 µg/ml poly-D-lysine (Gibco, A3890401). Cells tested negative for mycoplasma.

### Cell engineering

The DNA sequence of the G-Flamp1 cAMP biosensor ^15^ was synthesized (Twist Bioscience) and integrated into the pTwist Lenti CAG Puro expression vector. We transfected the expression vector using PEI, together with second-generation packaging plasmids into the Lenti-X 293T cell line (Takara Bio, 632180). Medium was changed 8 hours after transfection and the virus-containing supernatant was collected after 48 hours. We infected ASMCs at an MOI < 1 for 24 h, and used the cells for experiments after 3 days. We also introduced pCAG-GFlamp1 to WT HEK293, Δ*Q-GRK*, and Δ*Q-GRK* / Δ*β-arr1/2* knockout lines by lentivirus mediated transduction (MOI < 1) and stable polyclonal populations were selected. For biosensor experiments in HEK293T cells, we cloned the G-Flamp1 coding region under the control of the *Ef1a* promoter into a *piggyBac* transposase-compatible plasmid ^55^. We cotransfected the plasmid into cells with Lipofectamine LTX (ThermoFisher, 15338030) together with the *piggyBac* transposase plasmid to enable genome integration, and selected for a stable polyclonal population.

The DNA sequence of the Halo-ADRB2 transgene was synthesized (Twist Bioscience) and integrated in the pTwist Lenti CMV BSD expression vector. We produced lentivirus as described above, transduced HEK293T cells (MOI < 1), and selected for a stable polyclonal population.

### RNA sequencing and analysis

RNA was extracted from cells using the Quick-RNA Miniprep kit (Zymo Research, R1054) according to the manufacturer’s standard protocol. The RNA-seq libraries were prepared and sequenced by Azenta Life Sciences; polyadenylated RNA was enriched, the quality assessed by capillary electrophoresis, and libraries sequenced on a NovaSeq (Illumina) platform to generate 150 bp paired-end reads (2 × 150 bp), with a target depth of 30 million reads per sample.

Reads were aligned to the human reference genome (GRCh38, Ensembl release 106) using STAR v2.7.1a ^56^ with a splice junction overhang of 150 bp and excluding reads mapping to more than 20 genomic loci. Read counts were quantified using featureCounts ^57^ in paired-end mode with strand specificity disabled, from which transcripts per million (TPM) was computed.

### Ligand treatment regimes

To create oscillatory ligand regimes, we added isoproterenol (Tocris Bioscience, 1747) at 10× concentration followed by full media changes at 30 min intervals. For each experiment, we included a vehicle condition in which no ligand is added but media changes are made. For constant pretreatment, we added isoproterenol once after a 30 min delay. We additionally utilized forskolin (Cayman Chemical, 11018) in place of isoproterenol at a range of concentrations.

### Time-lapse biosensor fluorescence microscopy

All time-lapse biosensor experiments were performed on a Ti2 inverted microscope (Nikon), with a SOLA V-nIR light source (Lumencor), Zyla 4.2 PLUS sCMOS camera (Andor), 10× 0.3 NA objective (Nikon), and an EGFP filter cube (Chroma, 49002). The microscope was enclosed inside an environmental chamber (Okolab) with humidified 5 % CO_2_ flow and a temperature of 37 °C. We acquired images at intervals of 40 seconds.

### Biosensor image and time series analysis

To correct for movement of the field of view caused by adding and removing the ligand, we applied an image registration algorithm ^58^ to the image stacks, implemented as a plugin inside of Fiji. After registration, we used a Cellpose-Trackmate pipeline to segment and track cells over time. We used Cellpose’s Cytoplasm 3.0 model ^59^ to segment cells, setting the cell diameter parameter to be 37 µm for ASMCs and 20 µm for HEK293T. After segmentation, we used TrackMate 7 ^60^ with the Overlap algorithm to track cells, setting the minimum overlap to 30 %, and the overlap calculation to be precise. We implemented this Cellpose-Trackmate pipeline in the TrackMate Fiji plugin.

We processed and plotted time series using Python and the Matplotlib package. First, we removed tracks with a duration less than half of the experiment, and ignored isolated single frame artifacts caused by autofocus failure. With the remaining tracks, we corrected for fluctuations in background and sensor fluorescence caused by plate manipulations by background-correcting the fluorescence using a null field of view. Finally, we calculated the fluorescence change of the tracks as *ΔF / F_0_*, where *F* is the fluorescence signal in counts, *F_0_* is the fluorescence signal at time 0, and *ΔF* represents the change in fluorescence from time 0 (*F* - *F_0_*).

In addition to plotting the time series, we quantified alertness as the ratio of the peak response from the stress and priming ligand concentration. We extracted the peaks from the *ΔF / F_0_* signal, defined as the maximum value occurring within 10 frames of an increase in ligand concentration, and then calculated the ratio of these means. For the cAMP decay-rate comparison (Fig. S4, A and B), we averaged the single-cell traces within each experiment, aligned the mean traces to their peak, and fit the decay from the peak to 60 min after the peak with a single exponential with offset, *F*(*t*) = *A*·*e*^-*k*·*t*^ + *C*, where *t* is time after the peak, *A* is the decay amplitude, *C* is the plateau offset, and *k* is the decay rate constant. The model was fit by nonlinear least squares and *k* was plotted.

### β2-adrenergic receptor imaging and analysis

Prior to being used for experiments, we stained Halo-ADRB2 polyclonal cells with Halo ligand conjugated with JF635i, at a concentration of 50 nM, in cell culture media at 37 °C for 10 minutes. After 3 washes in Dulbecco’s phosphate buffered saline (DPBS), we trypsinized cells and sorted them by fluorescence activated cell sorting for cells with the top 15 % fluorescent intensity. We allowed these cells to recover for 6 days before use in experiments. During experiments, we observed the change in surface receptor fluorescence by pseudo-time lapse. We exposed cells in different wells of a plate to 3 μM isoproterenol in cell culture media for different durations between 30 and 120 minutes. In other wells, we exposed cells to isoproterenol for 120 minutes before washing with cell culture media and allowing cells to recover for durations between 30 and 150 minutes. Afterwards, we fixed cells for 10 minutes in 1 % paraformaldehyde, aligning all conditions in time so they were fixed at the same time. Next, we washed cells twice in DPBS, and stained them with JF549i-conjugated HaloTag ligand at a concentration of 50 nM, in DPBS at room temperature for 40 minutes. HaloTag ligands were a gift from Luke Lavis (HHMI Janelia). We acquired images on a Ti2 inverted microscope (Nikon), with a SOLA V-nIR light source (Lumencor), Zyla 4.2 PLUS sCMOS camera (Andor), 20× 0.5 NA objective (Nikon), and an ET Gold FISH filter cube (Chroma, 49304).

For analysis of surface membrane fluorescence, we segmented individual cells with Cellpose’s Cytoplasm 3.0 model ^59^, setting the cell diameter parameter to be 20 µm. Within each cell mask, we defined the plasma membrane as the 3-pixel-wide band at the cell perimeter and the remaining mask as the cell interior. For each cell, we computed a background-corrected surface signal as the median membrane-pixel intensity minus the median interior-pixel intensity; because internalized receptor is not labeled in this surface assay, the cell interior reports local autofluorescence and serves as a per-cell background. To exclude debris, we included only cells with more than 200 pixels in the membrane ring. Within each independent experiment, we normalized the median per-cell surface signal at each timepoint to the median at *t* = 0.

### Three-state receptor mathematical model

We employed a three-state receptor model that incorporates ligand binding, receptor desensitization, and receptor resensitization. Using the assumption of mass action kinetics, we modeled the transition of the receptor between the three states: free and unbound (*R_free_*), bound by ligand (*R_active_*), and desensitized (*R_desen_*). Transitions between *R_free_* and *R_active_* represent ligand– receptor association and dissociation at the cell membrane. The transition from *R_active_* to *R_desen_* represents receptor desensitization, while the transition from *R_desen_* to *R_free_* represents receptor resensitization. We modeled these processes phenomenologically to aggregate the effects of multiple molecular mechanisms regulating receptor sensitivity. The total receptor population is assumed to be conserved such that *R_free_(t)* + *R_active_(t)* + *R_desen_(t)* = *R_total_.* The dynamics of the receptor states are described by the following system of ordinary differential equations:

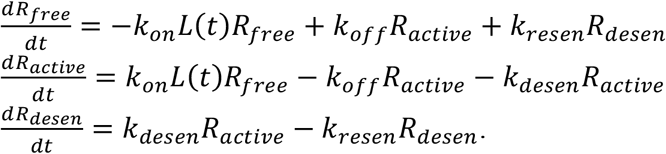

Here *k_on_* and *k_off_* denote the ligand–receptor association and dissociation rate constants, while *k_desen_* and *k_resen_* represent receptor desensitization and resensitization rate constants, respectively. *L* represents the concentration of the ligand. The downstream signaling output, cAMP, is assumed to be proportional to the active receptor population such that *cAMP*(*t*) = *R_active_*(*t*).

We varied the ligand concentration *L* with time to make a time course consisting of a pretreatment phase followed by a step change, with the pretreatment being either constant or oscillatory. Time *t* = 0 denotes the onset of ligand; both regimes are preceded by a ligand-free period of length *P*/2, where *P* is the oscillation period, which keeps the two conditions synchronized and during which the system rests at its ligand-free steady state. The pretreatment then proceeds for a duration *τ* at concentration *L_initial_*, delivered either continuously (constant pretreatment) or as a square wave oscillating between *L_initial_* and *L_trough_* with duty cycle *D* (oscillatory pretreatment). At time *τ*, *L* steps to a new constant concentration *L_final_* for the remainder of the simulation. Thus, in the constant pretreatment regime,

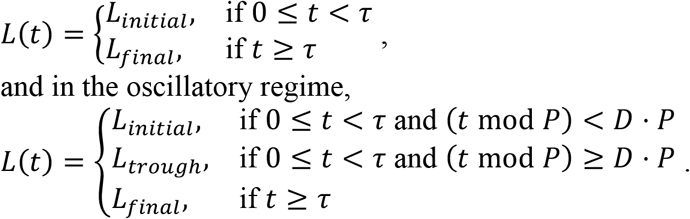

In most simulations we fixed *L_trough_* = 0, *D* = 0.5, and *L_initial_* = *K_d_*, and varied *L_final_* across 28 values spaced logarithmically from 10^-10^ to 10^-5^ M. In Fig. S6, we additionally varied *L_initial_*, *L_trough_*, and *D*.

Because the model is linear, we simulated it analytically. For any fixed ligand concentration, the three equations can be reduced to one matrix equation d**y**/d*t* = **A**(*L*) **y**, where **y**(*t*) is a column vector of the three receptor states (*R_free_*, *R_active_*, *R_desen_*) and **A**(*L*) is a 3×3 matrix of transition rates between receptor states: association, dissociation, desensitization, and resensitization. The ligand concentration modifies the rate matrix through the association rate. Because both ligand regimes hold the concentration constant in steps, the receptor states over each step can be found exactly by taking the exponential of the matrix, **y**(*t*) = exp[**A**(*L*)(*t* − *t_0_*)] **y**(*t_0_*). We evaluated this matrix exponential from the eigendecomposition of **A**(*L*) and carried the solution forward step by step, using the states at the end of one step as the starting values for the next. Simulations began with all receptors in the free state (*R_free_* = 1, *R_active_* = 0, *R_desen_* = 0), and cAMP was taken as proportional to *R_active_*.

### Extended receptor kinetic mathematical models

We introduced two extended versions of the model with increased molecular detail. In the first, which we refer to as the ‘cAMP turnover’ model, we included cAMP synthesis and degradation. We modeled cAMP synthesis and degradation as first order processes, such that the dynamics of cAMP are described by:

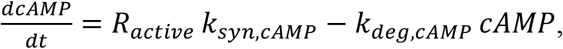

where *k_syn,cAMP_* is the rate constant for cAMP synthesis and *k_deg,cAMP_* is the rate constant for cAMP degradation. The starting conditions were *R_total_ =* 1*, R_free_* = *R_total_*, *R_active_* = 0, *R_desen_* = 0 and *cAMP* = 0. All other simulation parameters and ligand regimes were identical to those described for the base model.

In the second extended model which we refer to as the ‘receptor turnover’ model, we introduced synthesis and degradation of receptors. In this formulation, the total receptor population is no longer conserved; instead, free receptors are synthesized at a constant rate *v_syn,mem_*, representing constitutive receptor expression, and receptors in each state are subject to first-order degradation. Membrane-resident receptors (*R_free_* and *R_active_*) are degraded at rate *k_deg,mem_*, reflecting basal receptor turnover at the cell surface, while desensitized receptors (*R_desen_*) are degraded at rate *k_deg,int_*, reflecting the accelerated degradation of internalized receptors following endocytosis. The dynamics of the receptor states are described by:

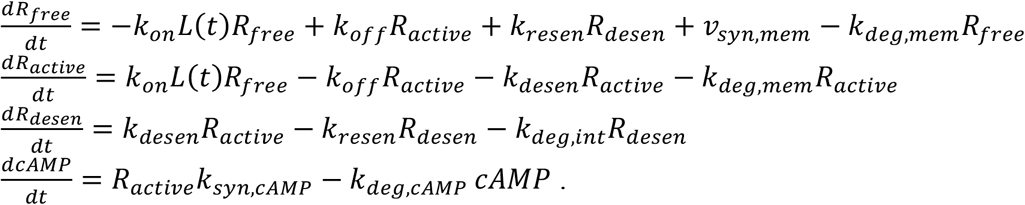

At steady state in the absence of ligand, the free receptor population is maintained at *R_free_* = *v_syn,mem_* / *k_deg,mem_*. We set the starting conditions to this steady-state *R_free_* value, with *R_active_* = 0, *R_desen_* = 0, and *cAMP* = 0. Ligand regimes were identical to those described for the base model.

Both alternative models were simulated analytically by the same matrix-exponential method used for the base model. The cAMP turnover model takes the form d**y**/d*t* = **A**(*L*)**y** as before, with the state vector **y** extended to include cAMP and **A**(*L*) a 4×4 rate matrix. The receptor turnover model uses the same four-component state vector but the constant synthesis term *v_syn,mem_* is zero-order giving d**y**/d*t* = **A**(*L*)**y** + **b**, where **b** places the synthesis rate in the free-receptor position and zeros elsewhere.

### Parameterization and sensitivity scan of the receptor models

The desensitization rate (*k_desen_*), resensitization rate (*k_resen_*), and the ligand binding rates (*k_on_* and *k_off_*) were estimated by fitting the base three-state receptor model to ASMC time course data during a step increase in isoproterenol concentration, from 60 nM to a saturating 3 μM. For each biological replicate we averaged the single-cell traces and scaled the mean trace to the maximum value under the matching naive condition. We then simulated this same step protocol for each replicate, matching its timing. Parameters were estimated by random search, drawing 20,000 parameter sets from log-uniform distributions within literature-based bounds (*k_desen_*, 10^-5^ to 10^-1^ s^-1^; *k_resen_*, 10^-6^ to 10^-2^ s^-1^; *k_on_*, 10^4^ to 10^8^ M^-1^ s^-1^; *k_off_*, 10^-3^ to 10^1^ s^-1^). For each set we scored the fit using three measures: the residuals between the simulated and measured time courses across the full trace, the residuals within a window around the second peak response, and the absolute deviation from the measured ratio of the second cAMP peak to the first. To combine these, we first divided each measure by its median across all sampled sets, which puts them on a common scale, and then added them together in a weighted sum that gave the window measure twice the weight of the other two. We selected the parameter set with the lowest combined score (table S2). For the HEK293 cells we estimated only the desensitization and resensitization rates in the same way, holding the binding rates fixed (table S3). For the alternative cAMP and receptor turnover models, we set the additional parameters using estimates from the literature (table S4).

To observe how sensitive alertness is to receptor desensitization specifically (Fig. 2I and J), we simulated the alertness protocol with the HEK293-fitted parameters, scaling *k_desen_* alone to 0.1×, 0.3×, or 1× its fitted value, while holding the other parameters at their optimal value.

### Comparison of oscillation with alternative strategies

For the low-ligand concentration strategy, we scanned the priming concentration from 2 to 49 % of steady-state receptor occupancy (25 linearly spaced values) with *R_total_* = 1, and set the post-shift concentration to 2× the priming occupancy fraction, so that the size of the stress step relative to baseline was held constant across the scan. For the high-receptor strategy, we used a priming concentration of 60 nM (EC_50_) and post-shift concentration of 3 µM (saturating), and scanned *R_total_* between 1 and 2 (17 values). The plotted representative low-ligand and high-receptor time series (Fig. 4C) used a priming concentration at EC_15_ and *R_total_* = 2×, respectively. For the low-ligand and high-receptor ‘combined’ condition, we varied operating concentration and *R_total_* to find parameters that matched oscillation on both alertness and tunability. We did this by first solving for the priming concentration that reproduced oscillation’s alertness via root-finding (Brent’s method), then scaled *R_total_* proportionally to match oscillations tunability. This gave a matched condition primed at EC_9_ with *R_total_* = 5.3×.

To test the robustness of the oscillatory and combined conditions, we performed a one-at-a-time parameter sensitivity analysis. Oscillation was centered at a priming concentration of 60 nM (EC_50_), and the combined condition was again matched by solving for the priming concentration that reproduces the oscillatory alertness, then scaling receptor abundance to reproduce its tunability. We first scanned the operating dose, drawing 1,000 lognormal multipliers with a geometric standard deviation of 1.5 and applying the same draws to both conditions. We fixed the fold change at 2, and the multiplier rescaled both the priming and stress dose. We next scanned *k_desen_* and *k_resen_* each with a geometric standard deviation of 3 and holding the other parameter at its center value. For the representative traces, we centered oscillation at 60 nM (EC_50_) and the combined condition at 6.3 nM (EC_9_), stepping each to twice its own occupancy, 3 µM (saturating) and 14 nM (EC_18_) respectively. We simulated four cells per condition, at *k_desen_* = 0.3, 1, 3, and 10 times the reference.

For every condition we computed alertness as previously: the ratio of the post-shift to the priming peak of cAMP. Because the low-concentration strategy is, by definition, restricted to a lower range of concentrations, we scaled the tunability metric’s range to each scenarios own priming concentration, rather than using one fixed dose for every scenario. Tunability was defined the largest post-shift peak response across this range: 21 log-spaced doses from 0.13× to 2× the priming occupancy fraction.

### Receptor availability across desensitization and resensitization rates

In Fig. 5A, we asked how oscillatory versus constant pretreatment changes the receptor species prior to a concentration step. Under constant ligand *L* the receptor steady state is the solution of **A**(*L*) **y** = 0, which we solved under receptor conservation (*R_free_* + *R_active_* + *R_desen_* = *R_total_*). Under oscillation, the receptor state is advanced over one period *P* by the matrix **M** = exp[**A**(*L*_trough_) *P*/2] exp[**A**(*L*_peak_) *P*/2]. The periodic steady state **y**\* is the state that is unchanged after one *P*, **M y**\* = **y**\*, which we solved under receptor conservation. Sweeping *k_desen_* and *k_resen_* over a 120×120 logarithmic grid (10^-5^ to 10^-1^ and 10^-6^ to 10^-2^ s^-1^), we read the available receptor pool (*R_free_* + *R_active_*) at the end of the off-phase, immediately before the step, and compared it between the oscillatory and constant pretreatment conditions.

### Rate-constant compilation across cell types

We assembled rate constants for functional desensitization and resensitization of the β-adrenergic receptor across cell types through a literature search (tables S5 and S6). We kept the focus deliberately functional: we excluded studies whose only readout was receptor-level (for example radioligand binding or receptor internalization assays) rather than a functional signaling output. More specifically, we included an entry only when it met all four conditions: (1) the receptor was a β-adrenergic receptor (β1, β2, or mixed) and the agonist was epinephrine, isoproterenol, or norepinephrine; (2) the readout reported cAMP accumulation or adenylyl cyclase activity; (3) desensitization or resensitization was induced in live cells; and (4) the study reported a time course. We identified studies through a guided search of PubMed; this was not a systematic review and qualifying studies may exist that these queries did not surface. Our queries combined receptor terms (‘β-adrenergic’, ‘beta-adrenergic’, ‘β-adrenoceptor’, and the subtype variants β1 and β2) and agonist terms (isoproterenol, isoprenaline, norepinephrine, catecholamine) with process terms (‘desensitization’, ‘resensitization’, ‘recovery’, ‘recovery of responsiveness’, ‘reversal of desensitization’) and functional-readout terms (‘adenylate/adenylyl cyclase’, ‘cAMP’). We followed the reference lists of retrieved articles to locate time-course studies. Where authors reported a rate constant, we used it directly; where they reported a half-time, we converted it according to *k* = ln 2 / *t_1/2_*. Otherwise we digitized the published time course with WebPlotDigitizer^61^ and fit a single exponential with offset, *y*(*t*) = *A*·*e*^-*kt*^ + *C* for desensitization and *y*(*t*) = *A*·(*1* - *e*^-*kt*^) + *C* for recovery, where *y* is the functional response (cAMP accumulation or adenylyl cyclase activity), *A* the amplitude, *k* the rate constant, and *C* the offset.

### Statistics

We performed all hypothesis tests on independent experiments, on per-experiment means. Comparisons between two conditions measured within the same experiments (constant vs. naive or constant vs. oscillation) used a paired two-tailed *t* test. Comparisons across genotypes used one-way ANOVA followed by Tukey HSD (Fig. 2G) or Welch’s ANOVA with Games-Howell when group variances were unequal (Fig. S4B). Dose-response curves were fitted with a four-parameter logistic model, with EC_50_ reported as the fit estimate and its 95 % CI.

---

**Fig. S1.**
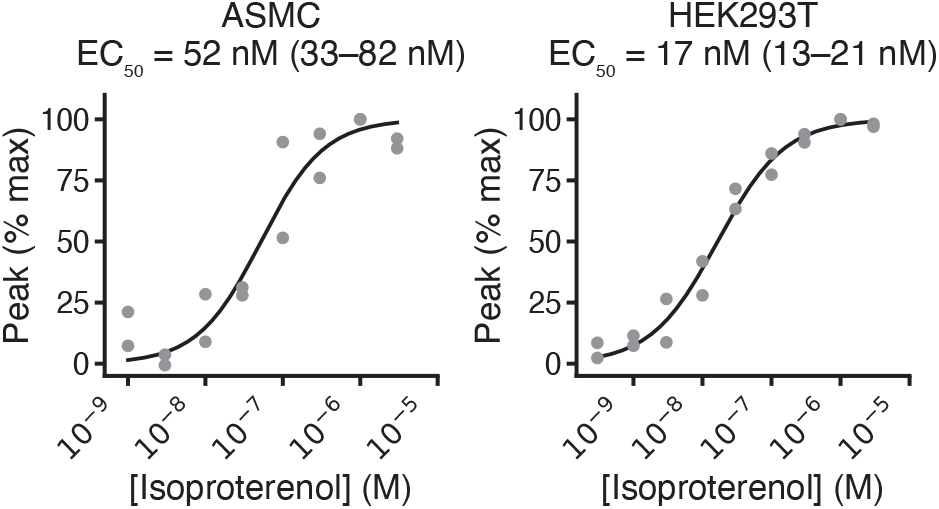
Dose-dependent activation of the cAMP biosensor by isoproterenol. Isoproterenol activates the cAMP biosensor in a dose-dependent manner in ASMCs and HEK293T cells. Points, per-experiment means normalized to maximum response; *n* = 2 independent experiments per dose. Curves, four-parameter logistic fits; EC_50_ shown with the 95 % CI of the fit.

**Fig. S2.**
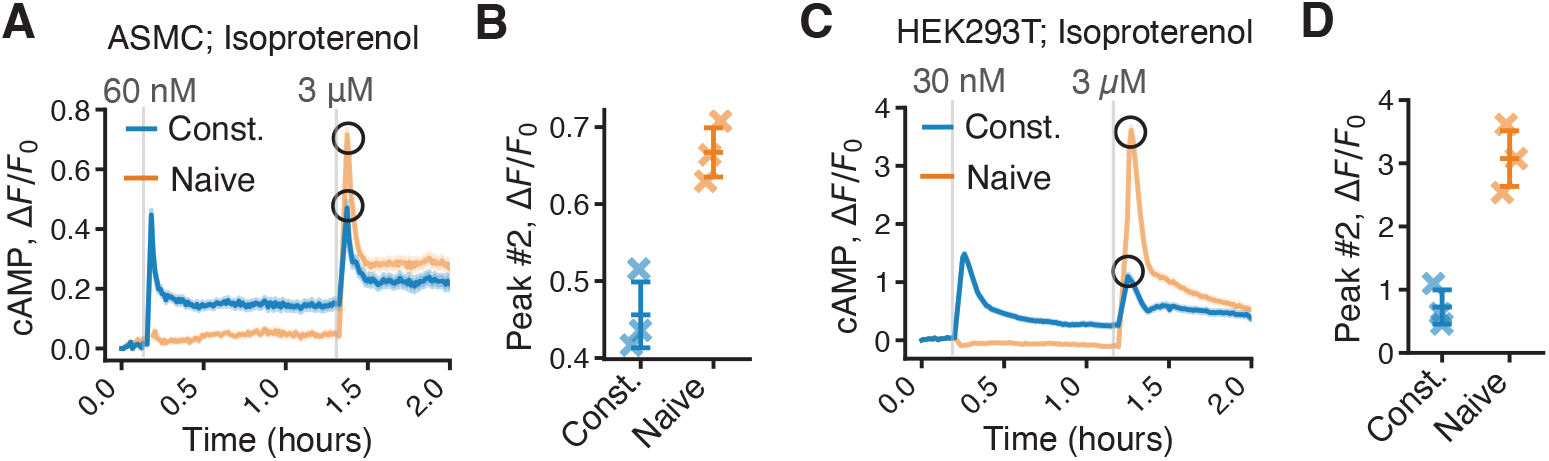
Cells lose alertness to changes in ligand concentration under constant pretreatment. (**A** and **C**) Representative cAMP time courses during the alertness protocol in ASMCs (A) or HEK293T cells (C), under constant pretreatment or naive conditions. Traces show the mean ± SEM for *n* = 370 (constant) and 428 (naive) cells in ASMC, and *n* = 910 (constant) and 511 (naive) cells in HEK293T. (**B** and **D**) Second-peak height (the response to 3 μM isoproterenol) under constant pretreatment and naive conditions, in ASMCs (B) and HEK293T cells (D). Crosses, individual experiments; bars, mean ± SD; *n* = 3 independent experiments for each cell type. Constant vs. naive, paired two-tailed *t* test: ASMC (B), *P* = 0.0073; HEK293T (D), *P* = 0.0034.

**Fig. S3.**
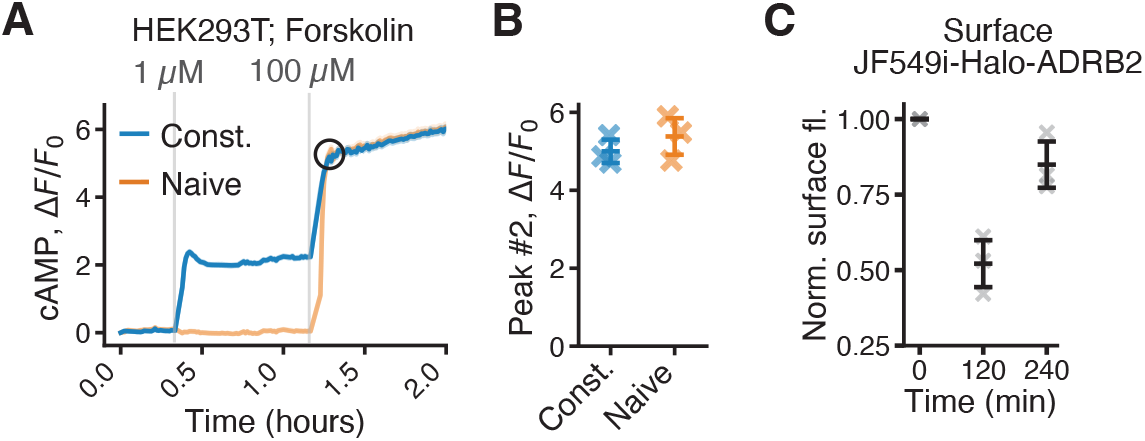
Loss of alertness originates at the receptor. (**A**) Representative cAMP time course during the alertness protocol in HEK293T cells stimulated with forskolin, under constant pretreatment or naive condition. Traces show the mean ± SEM for *n* = 737 (constant) and 262 (naive) cells. (**B**) Second-peak height (the response to 100 μM forskolin) under constant pretreatment and naive conditions. Crosses, individual experiments; bars, mean ± SD; *n* = 3 independent experiments. Constant vs. naive, paired two-tailed *t* test: *P* = 0.45. (**C**) Quantification of normalized surface fluorescence intensity (JF549i-Halo-ADRB2) in HEK293T cells during isoproterenol treatment (0–120 min) and washout (120–240 min), showing surface receptor loss and subsequent partial recovery. Values are normalized per experiment to the *t* = 0 baseline. Gray crosses, individual experiments; bars, mean ± SD; *n* = 3 independent experiments. Loss (0 vs. 120 min), paired two-tailed *t* test: *P* = 0.013; recovery (240 vs. 120 min): *P* = 0.087.

**Fig. S4.**
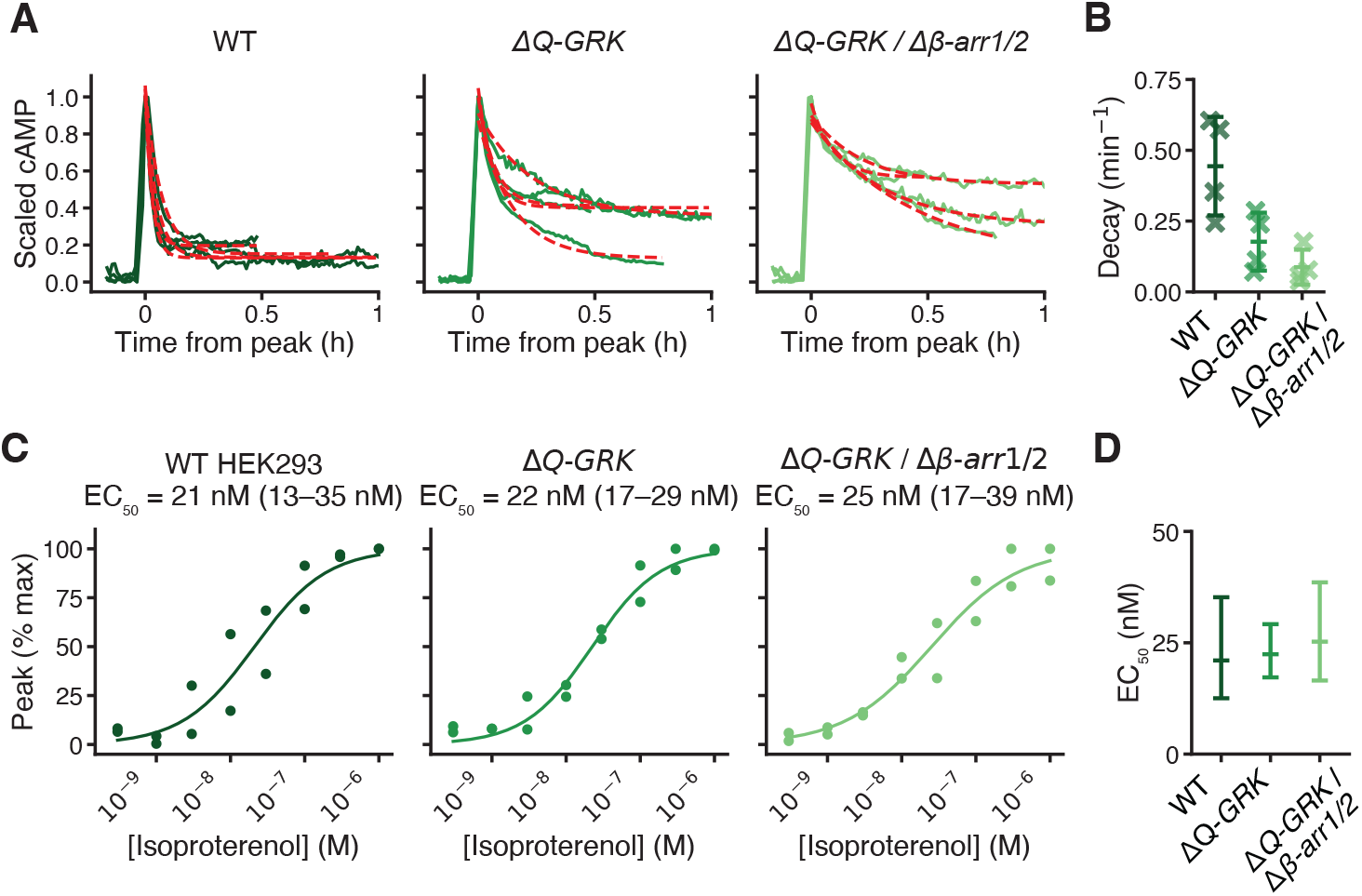
Loss of GRKs and *β-arrestin 1/2* slows apparent cAMP decay independent of potency. (**A**) Scaled cAMP time courses following the response to isoproterenol in WT, Δ*Q-GRK*, and Δ*Q-GRK* / *Δβ-arr1/2* HEK293 cells, aligned to the peak (*t* = 0). Each colored trace is the mean across cells from one independent experiment (*n* = 4 independent experiments), with the fitted decay model overlaid (red dashed). (**B**) Quantification of cAMP decay rate following the peak response to isoproterenol in WT, Δ*Q-GRK*, and Δ*Q-GRK* / Δ*β-arr1/2* HEK293 cells. Crosses, individual experiments; bars, mean ± SD; *n* = 4 independent experiments. Welch’s ANOVA: *P* = 0.033. Games-Howell *post hoc* vs. WT: Δ*Q-GRK*, *P* = 0.10; Δ*Q-GRK* / Δ*β-arr1/2*, *P* = 0.043. (**C**) Dose-response curves of peak cAMP in WT HEK293, Δ*Q-GRK*, and Δ*Q-GRK* / Δ*β-arr1/2*. Points, per-experiment means normalized to the maximum response; *n* = 2 independent experiments per dose. Curves, four-parameter logistic fits; EC_50_ shown with the 95% CI of the fit. (**D**) EC₅₀ values from the fits in (C). Bars, fitted EC₅₀ with 95% CI.

**Fig. S5.**
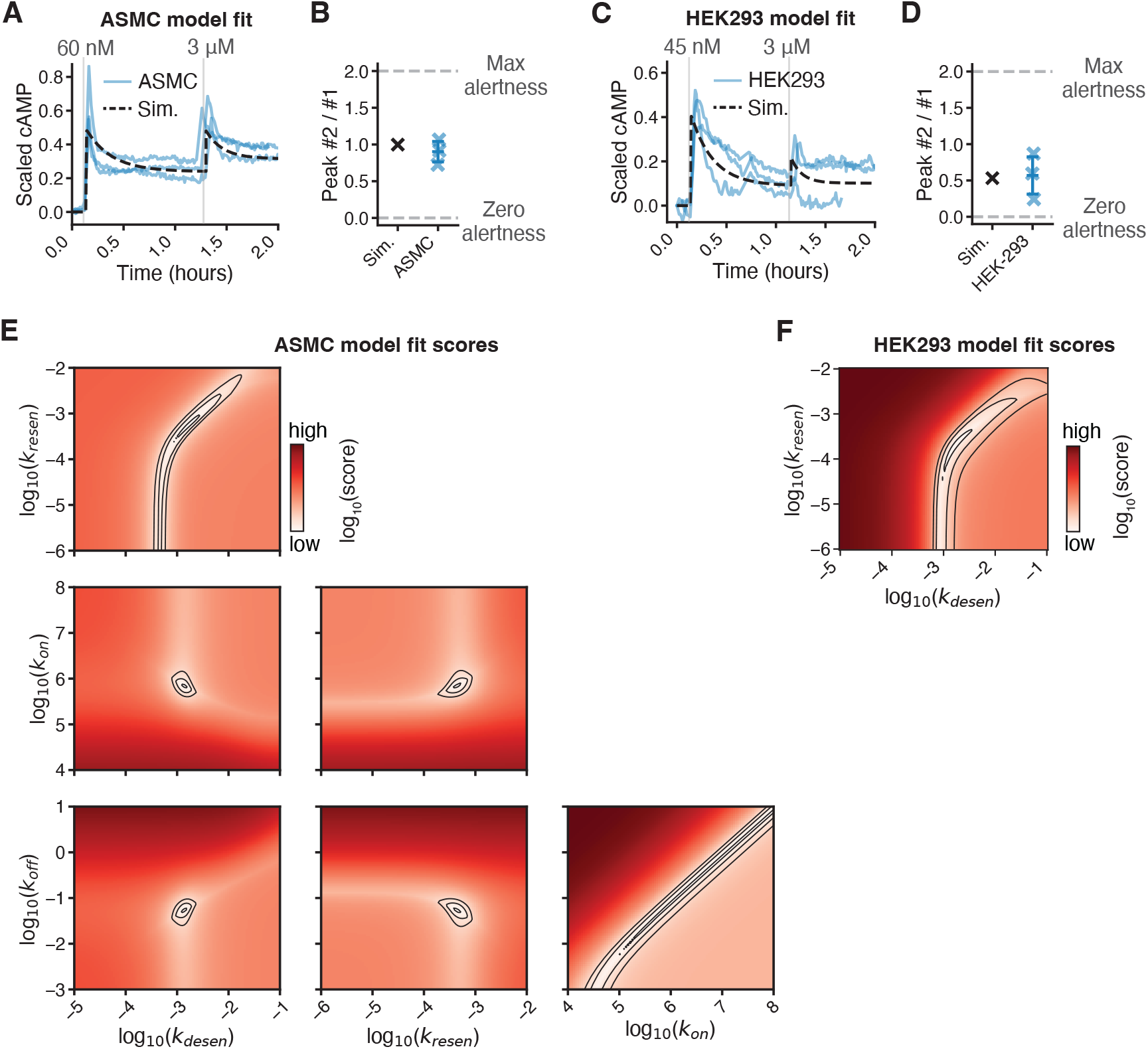
The three-state receptor model captures the alertness of ASMCs and HEK293 cells. (**A**) Scaled cAMP time courses during the alertness protocol in ASMCs; each light blue trace is the mean of one independent experiment (*n* = 3 independent experiments), with the fitted model simulation overlaid. (**B**) The ratio of peak heights (peak #2 / #1) in (A), for ASMCs and the model simulation. (**C** and **D**) As in (A and B), for HEK293 cells during the alertness protocol. (**E** and **F**) We varied the parameters of the model pairwise, keeping the remaining parameters at the optimum value, and scored the fit to ASMC data (E) or HEK293 (F) time-course data. Low scores (white) indicate a better fit (see Methods for score details). Black contours are drawn at 1.1, 1.5 and 2× the lowest score. The model was fit sequentially: first, all four parameters were estimated from ASMC data (E); the resulting binding rates (*k_on_* and *k_off_*) were then held fixed while only the desensitization and resensitization rates were re-estimated from HEK293 data (F).

**Fig. S6.**
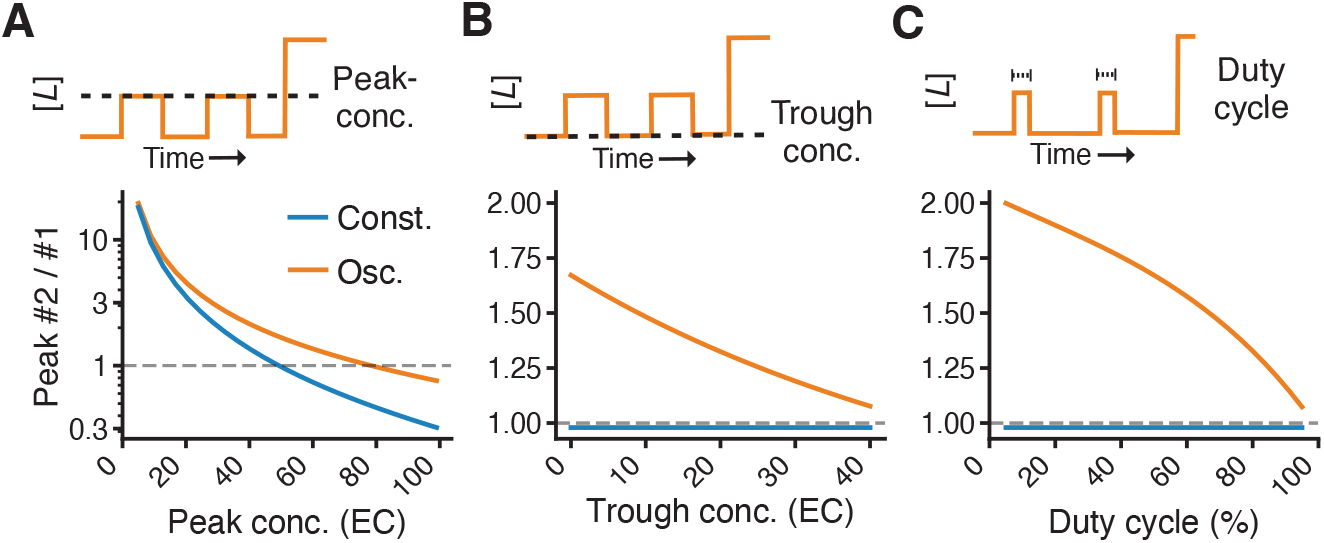
The dependence of peak response ratios on oscillation parameters. (**A** to **C**) The ratio of the peak heights (peak #2 / #1) under the alertness protocol, as a function of the peak concentration (A), trough concentration (B), or duty cycle of the pretreatment phase (C).

**Fig. S7.**
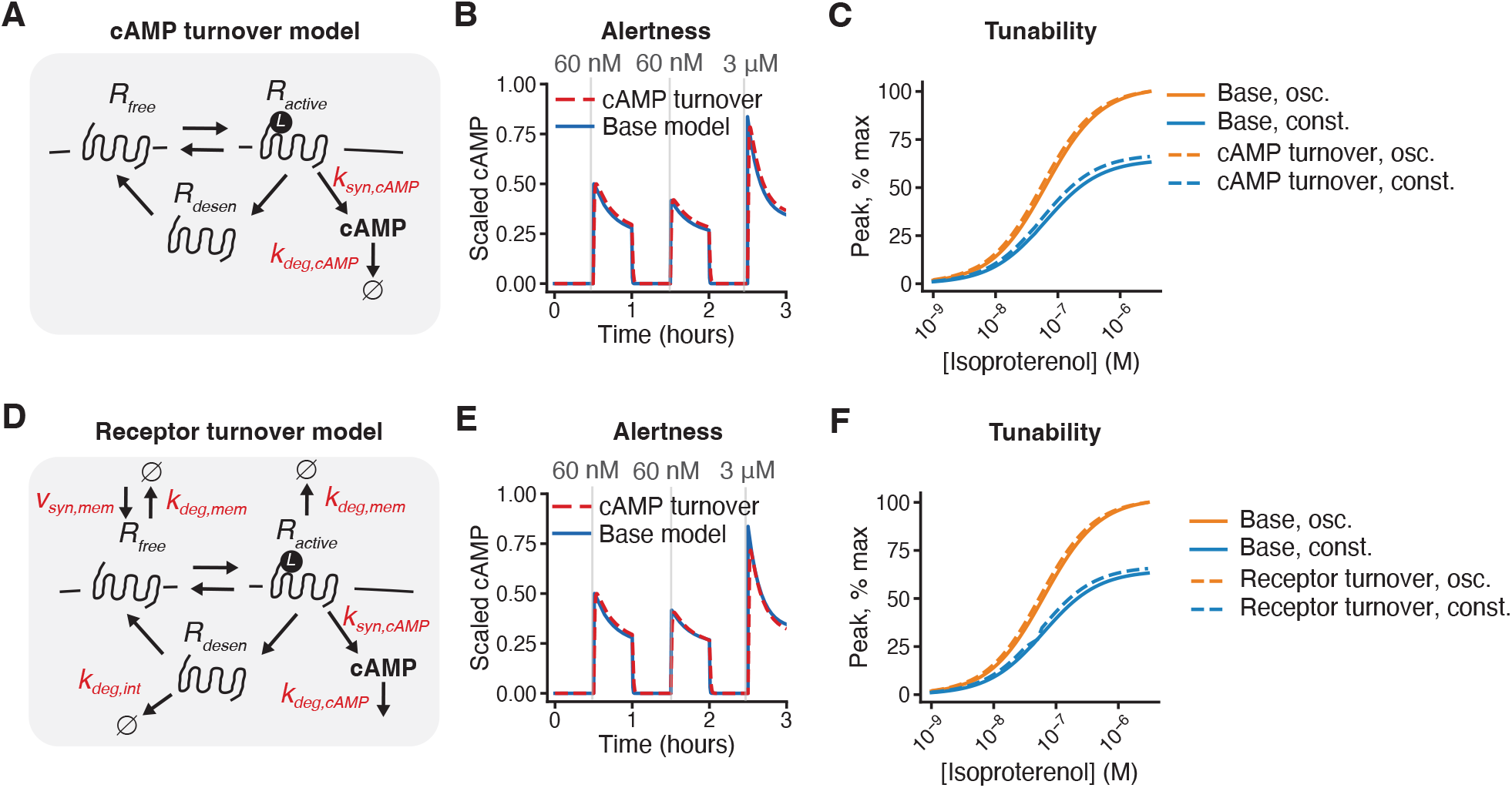
Extended kinetic models of β-adrenergic signaling show similar loss of alertness under constant pretreatment. (**A**) cAMP turnover model: we extend the base three-state receptor model with cAMP synthesis downstream of the active receptor (*k_syn,cAMP_*) and cAMP degradation (*k_deg,cAMP_*). (**B**) Simulated cAMP time courses for the alertness protocol in the base and cAMP turnover models. (**C**) Simulated dose-response curves of peak cAMP under the tunability protocol, for the base and cAMP turnover models, each under naive or constant pretreatment. (**D**) Receptor turnover model: we further extend (A) with membrane receptor synthesis (*v_syn,mem_*) and degradation of the membrane (*k_deg,mem_*) and desensitized receptor (*k_deg,int_*). (**E**) As in (B), for the receptor turnover model. (**F**) As in (C), for the receptor turnover model.

**Fig. S8.**
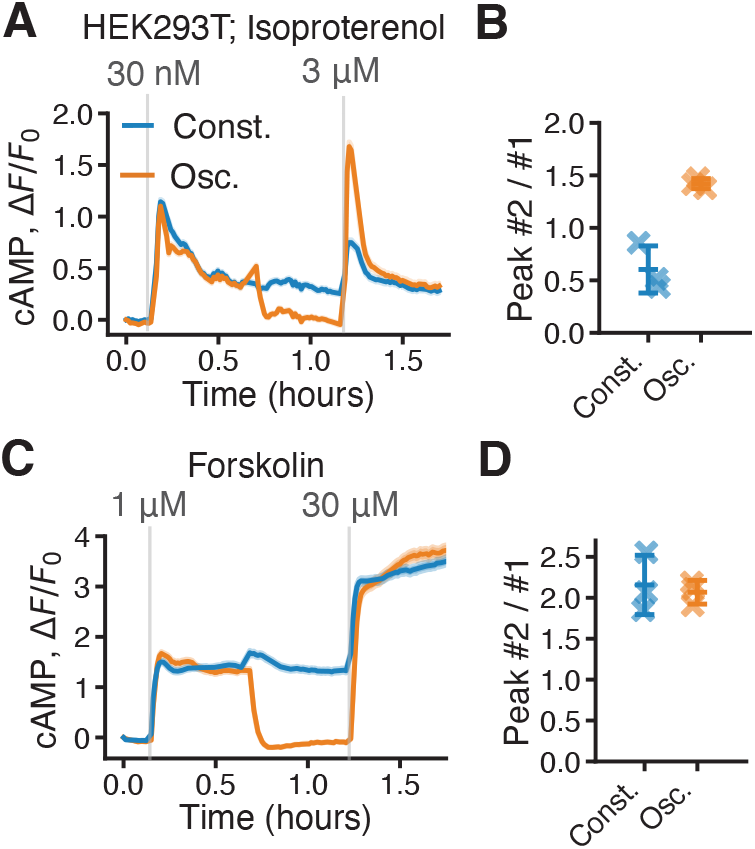
Oscillation improves alertness to isoproterenol but not forskolin in HEK293T cells. (**A** and **C**) Representative cAMP time courses during the alertness protocol, for isoproterenol (A) or forskolin (C), under constant or oscillatory pretreatment. Traces show the mean ± SEM for *n* = 611 (constant) and 774 (oscillatory) cells in (A), and *n* = 509 (constant) and 466 (oscillatory) cells in (C). (**B** and **D**) The ratio of peak heights (peak #2 / #1) for isoproterenol (B) or forskolin (D). Crosses, individual experiments; bars, mean ± SD; *n* = 3 independent experiments. Constant vs. oscillation, Paired two-tailed *t* test: isoproterenol (B), *P* = 0.025; forskolin (D), *P* = 0.74.

**Fig. S9.**
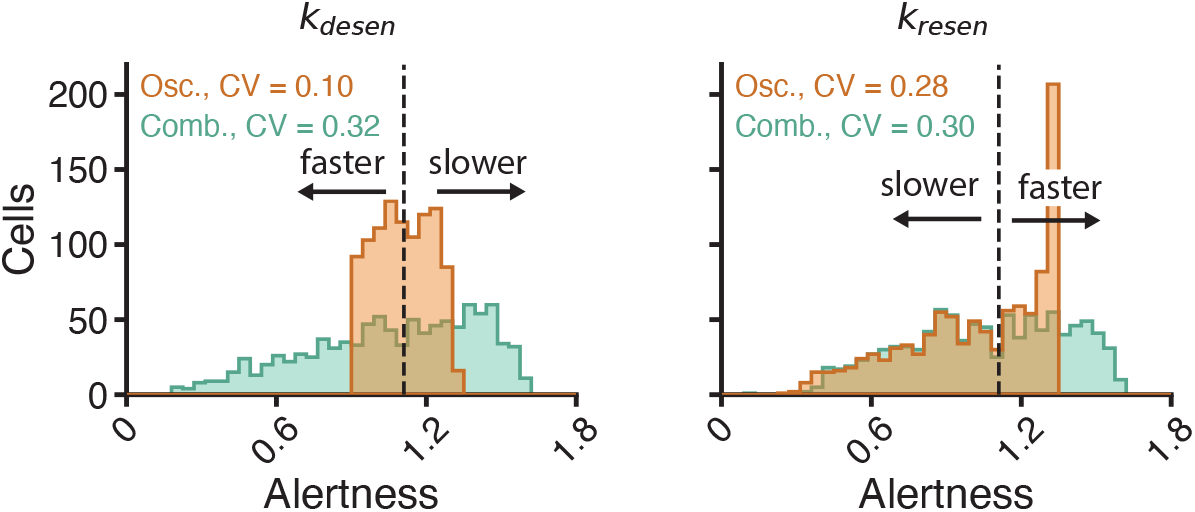
Oscillation improves consistency of the response to stress across receptor kinetics. Distributions of the simulated alertness following oscillatory or combined constant pretreatment, across 1,000 parameter sets in which *k_desen_* (*left*) or *k_resen_* (*right*) was sampled.

**Table S1.** β-adrenergic receptor subtype mRNA expression in ASMC and HEK293T.

| Gene | ASMC (TPM) | HEK293T (TPM) |
| --- | --- | --- |
| *ADRB1* | 0.0 | 0.8 |
| *ADRB2* | 0.9 | 2.2 |
| *ADRB3* | 0.0 | 0.0 |

TPM calculated from one sample per cell type.

**Table S2.**
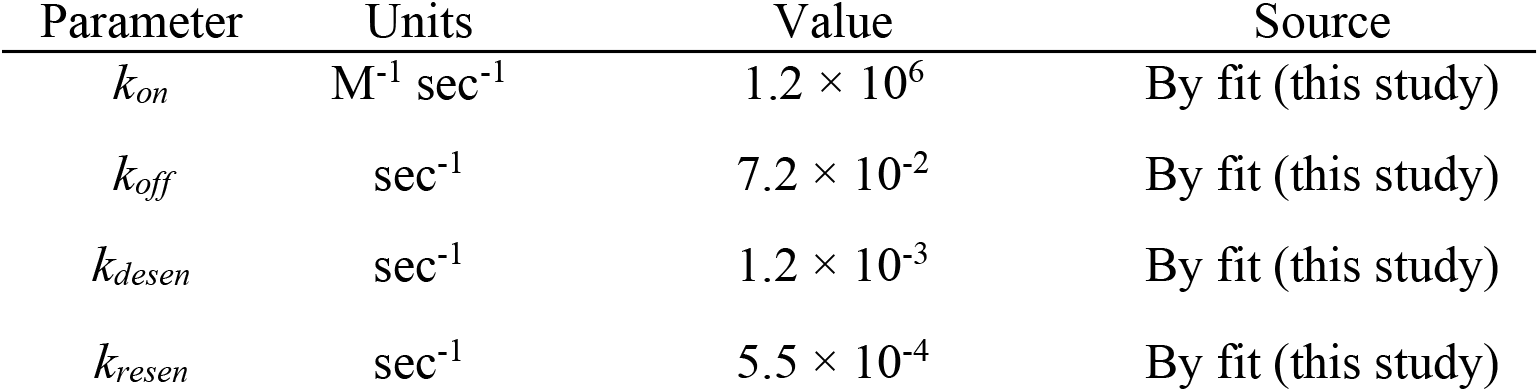
The parameter values used for the ASMC three-state receptor model.

**Table S3.** The parameter values used for the HEK293 three-state receptor model.

| Parameter | Units | Value | Source |
| --- | --- | --- | --- |
| $k_{on}$ | $M^{-1} sec^{-1}$ | $1.2 \times 10^6$ | By fit (this study) |
| $k_{off}$ | $sec^{-1}$ | $7.2 \times 10^{-2}$ | By fit (this study) |
| $k_{desen}$ | $sec^{-1}$ | $2.3 \times 10^{-3}$ | By fit (this study) |
| $k_{resen}$ | $sec^{-1}$ | $2.6 \times 10^{-4}$ | By fit (this study) |

**Table S4.**
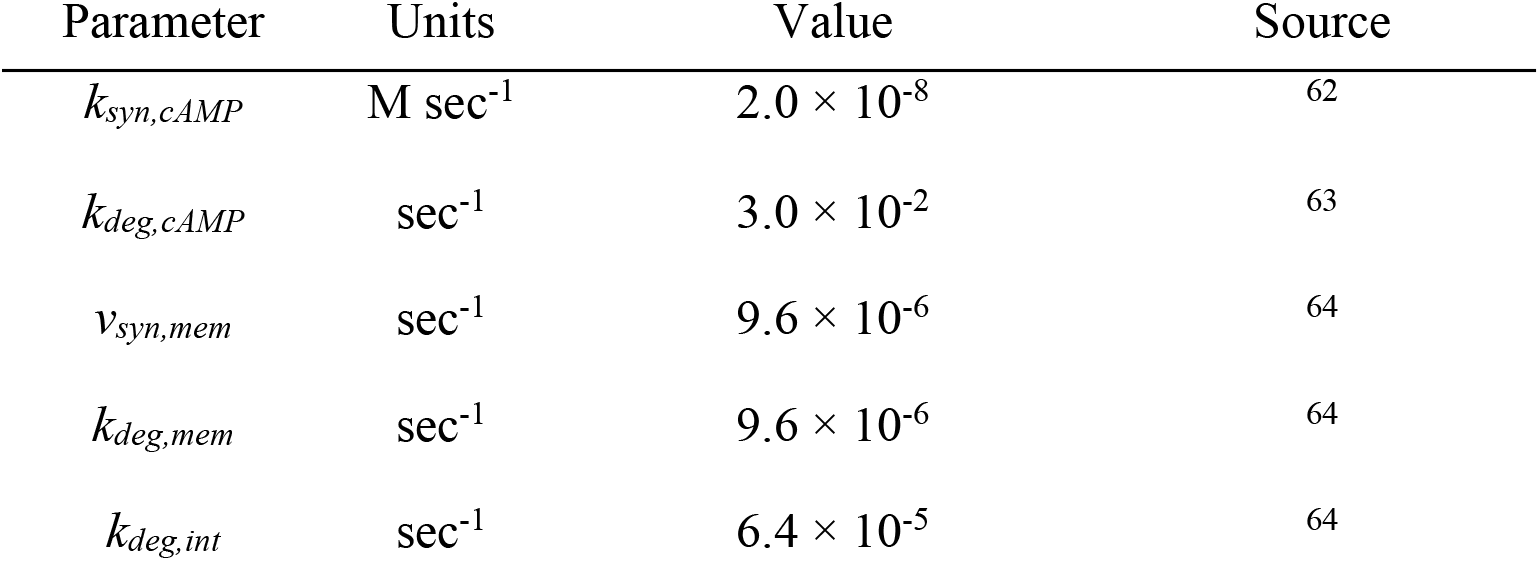
The additional parameter values used for the alternative receptor models.

**Table S5.** Observed signaling decay rates across cell types.

| Cell type | Receptor subtype | Agonist | Decay rate (sec<sup>-1</sup>) | Source |
| --- | --- | --- | --- | --- |
| HEK293* | β2 | epinephrine | $3.0 \times 10^{-2}$ | 65 |
| HEK293 | β2 | isoproterenol | $1.2 \times 10^{-2}$ | 38 |
| HEK293 | β2 | isoproterenol | $6.3\text{--}8.3 \times 10^{-3}$ | 66 |
| Chinese hamster fibroblast (CHW)* | β1 | isoproterenol | $4.3 \times 10^{-3}$ | 67 |
| 1321N1 astrocytoma | β<sup>†</sup> | norepinephrine | $1.2 \times 10^{-3}$ | 68 |
| Chick ventricular myocytes | β1/β2 | isoproterenol | $8.6 \times 10^{-4}$ | 69 |
| Frog erythrocyte | β1 | isoproterenol | $4 \times 10^{-4}$ | 18 |
| Turkey erythrocyte | β1 | isoproterenol | $3.9 \times 10^{-4}$ | 70 |
| Rat cortical astrocytes | β<sup>†</sup> | norepinephrine | $2.6 \times 10^{-4}$ | 71 |
| Rat hepatocytes | β<sup>†</sup> | isoproterenol | $2.3 \times 10^{-3}$ | 72 |
| Human neutrophil | β2 | isoproterenol | $2.7 \times 10^{-3}$ | 73 |
| Rat myometrium | β<sup>†</sup> | isoproterenol | $1.4 \times 10^{-3}$ | 74 |

\*The receptor was expressed transgenically. <sup>†</sup>The β subtype was not resolved.

**Table S6.** Observed signaling resensitization rates across cell types.

| Cell type | Receptor subtype | Agonist | Resensitization rate (sec<sup>-1</sup>) | Source |
| --- | --- | --- | --- | --- |
| Chinese hamster ovary (CHO)* | β2 | isoproterenol | $4 \times 10^{-3}$ | 75 |
| Striatal neurons | β<sup>†</sup> | isoproterenol | $1.6 \times 10^{-3}$ | 76 |
| Cortical neurons | β<sup>†</sup> | isoproterenol | $1.4 \times 10^{-3}$ | 76 |
| Chick ventricular myocytes | β1/β2 | isoproterenol | $7.4 \times 10^{-4}$ | 69 |
| Turkey erythrocyte | β1 | isoproterenol | $6 \times 10^{-5}$ | 70 |
| Rat hepatocytes | β<sup>†</sup> | isoproterenol | $7.0 \times 10^{-4}$ | 72 |
| Human neutrophil | β2 | isoproterenol | $1.4 \times 10^{-4}$ | 73 |
| Rat myometrium | β<sup>†</sup> | isoproterenol | $8.6 \times 10^{-5}$ | 74 |

\*The receptor was expressed transgenically. <sup>†</sup>The β subtype was not resolved.

**Movie S1. ASMCs adapted to isoproterenol lose alertness to a subsequent increase in concentration.**

Live-cell biosensor fluorescence imaging of cAMP in ASMCs. *Left*, cells pretreated with 60 nM isoproterenol before a step to 3 µM (Const.). *Right*, cells exposed directly to 3 µM isoproterenol (Naive). Time, min:s; agonist added at 37:37 and 109:10; scale bar, 200 µm. Representative of *n* = 3 independent experiments.

**Movie S2. Loss of alertness is reproduced in HEK293T cells.**

Live-cell biosensor imaging of cAMP in HEK293T cells. *Left*, cells preincubated with 30 nM isoproterenol before a step to 3 µM (Const.). *Right*, cells exposed directly to 3 µM isoproterenol (Naive). Time, min:s; agonist added at 37:10 and 97:57; scale bar, 200 µm. Representative of *n* = 3 independent experiments.

## References

1. Rajagopal, S., and Shenoy, S.K. (2018). GPCR desensitization: acute and prolonged phases. Cell. Signal. 41, 9–16.

2. Boucher, J., Kleinridders, A., and Kahn, C.R. (2014). Insulin receptor signaling in normal and insulin-resistant states. Cold Spring Harb. Perspect. Biol. 6, a009191. 10.1101/cshperspect.a009191.

3. Oakley, R.H., and Cidlowski, J.A. (1993). Homologous down regulation of the glucocorticoid receptor: the molecular machinery. Crit. Rev. Eukaryot. Gene Expr. 3, 63–88.

4. Boluyt, M.O., Long, X., Eschenhagen, T., Mende, U., Schmitz, W., Crow, M.T., and Lakatta, E.G. (1995). Isoproterenol infusion induces alterations in expression of hypertrophy-associated genes in rat heart. Am. J. Physiol. 269, H638–647.

5. Engelhardt, S., Hein, L., Wiesmann, F., and Lohse, M.J. (1999). Progressive hypertrophy and heart failure in β1-adrenergic receptor transgenic mice. Proc. Natl. Acad. Sci. U. S. A. 96, 7059–7064. 10.1073/pnas.96.12.7059.

6. Liggett, S.B., Tepe, N.M., Lorenz, J.N., Canning, A.M., Jantz, T.D., Mitarai, S., Yatani, A., and Dorn, G.W. (2000). Early and delayed consequences of β2-adrenergic receptor overexpression in mouse hearts: critical role for expression level. Circulation 101, 1707–1714. 10.1161/01.cir.101.14.1707.

7. Kudej, R.K., Iwase, M., Uechi, M., Vatner, D.E., Oka, N., Ishikawa, Y., Shannon, R.P., Bishop, S.P., and Vatner, S.F. (1997). Effects of chronic β-adrenergic receptor stimulation in mice. J. Mol. Cell. Cardiol. 29, 2735– 2746. 10.1006/jmcc.1997.0508.

8. Teerlink, J.R., Pfeffer, J.M., and Pfeffer, M.A. (1994). Progressive ventricular remodeling in response to diffuse isoproterenol-induced myocardial necrosis in rats. Circ. Res. 75, 105–113. 10.1161/01.res.75.1.105.

9. Noma, T., Lemaire, A., Naga Prasad, S.V., Barki-Harrington, L., Tilley, D.G., Chen, J., Le Corvoisier, P., Violin, J.D., Wei, H., Lefkowitz, R.J., et al. (2007). β-Arrestin–mediated β1-adrenergic receptor transactivation of the EGFR confers cardioprotection. J. Clin. Invest. 117, 2445–2458. 10.1172/JCI31901.

10. Oliver, G., and Schäfer, E.A. (1895). The physiological effects of extracts of the suprarenal capsules. J. Physiol. 18, 230–276. 10.1113/jphysiol.1895.sp000564.

11. Cannon, W.B., Shohl, A.T., and Wright, W.S. (1911). Emotional glycosuria. Am. J. Physiol. 29, 280–287. 10.1152/ajplegacy.1911.29.2.280.

12. Solis-Cohen, S. (1900). The use of adrenal substance in the treatment of asthma. J. Am. Med. Assoc. 34, 1164–1166. 10.1001/jama.1900.24610190014001c.

13. Warren, J.B., Dalton, N., Turner, C., and Clark, T.J.H. (1984). Protective effect of circulating epinephrine within the physiologic range on the airway response to inhaled histamine in nonasthmatic subjects. J. Allergy Clin. Immunol. 74, 683–686. 10.1016/0091-6749(84)90230-6.

14. Carstairs, J.R., Nimmo, A.J., and Barnes, P.J. (1985). Autoradiographic visualization of β-adrenoceptor subtypes in human lung. Am. Rev. Respir. Dis. 132, 541–547. 10.1164/arrd.1985.132.3.541.

15. Wang, L., Wu, C., Peng, W., Zhou, Z., Zeng, J., Li, X., Yang, Y., Yu, S., Zou, Y., Huang, M., et al. (2022). A high-performance genetically encoded fluorescent indicator for in vivo cAMP imaging. Nat. Commun. 13, 5363. 10.1038/s41467-022-32994-7.

16. Chuang, D.M., and Costa, E. (1979). Evidence for internalization of the recognition site of β-adrenergic receptors during receptor subsensitivity induced by (-)-isoproterenol. Proc. Natl. Acad. Sci. U. S. A. 76, 3024–3028. 10.1073/pnas.76.6.3024.

17. Stadel, J.M., Nambi, P., Shorr, R.G., Sawyer, D.F., Caron, M.G., and Lefkowitz, R.J. (1983). Catecholamine-induced desensitization of turkey erythrocyte adenylate cyclase is associated with phosphorylation of the β-adrenergic receptor. Proc. Natl. Acad. Sci. U. S. A. 80, 3173–3177. 10.1073/pnas.80.11.3173.

18. Sibley, D.R., Strasser, R.H., Benovic, J.L., Daniel, K., and Lefkowitz, R.J. (1986). Phosphorylation/dephosphorylation of the β-adrenergic receptor regulates its functional coupling to adenylate cyclase and subcellular distribution. Proc. Natl. Acad. Sci. U. S. A. 83, 9408–9412.

19. Ferguson, S.S., Downey, W.E., Colapietro, A.M., Barak, L.S., Ménard, L., and Caron, M.G. (1996). Role of β-arrestin in mediating agonist-promoted G protein-coupled receptor internalization. Science 271, 363–366. 10.1126/science.271.5247.363.

20. Benovic, J.L., Strasser, R.H., Caron, M.G., and Lefkowitz, R.J. (1986). β-adrenergic receptor kinase: identification of a novel protein kinase that phosphorylates the agonist-occupied form of the receptor. Proc. Natl. Acad. Sci. U. S. A. 83, 2797–2801. 10.1073/pnas.83.9.2797.

21. Lohse, M.J., Benovic, J.L., Codina, J., Caron, M.G., and Lefkowitz, R.J. (1990). β-Arrestin: a protein that regulates β-adrenergic receptor function. Science 248, 1547–1550. 10.1126/science.2163110.

22. Torphy, T.J., Zhou, H.L., and Cieslinski, L.B. (1992). Stimulation of β adrenoceptors in a human monocyte cell line (U937) up-regulates cyclic AMP-specific phosphodiesterase activity. J. Pharmacol. Exp. Ther. 263, 1195– 1205.

23. Manning, C.D., McLaughlin, M.M., Livi, G.P., Cieslinski, L.B., Torphy, T.J., and Barnette, M.S. (1996). Prolonged β adrenoceptor stimulation up-regulates cAMP phosphodiesterase activity in human monocytes by increasing mRNA and protein for phosphodiesterases 4A and 4B. J. Pharmacol. Exp. Ther. 276, 810–818.

24. Noda, C., Shinjyo, F., Tomomura, A., Kato, S., Nakamura, T., and Ichihara, A. (1984). Mechanism of heterologous desensitization of the adenylate cyclase system by glucagon in primary cultures of adult rat hepatocytes. J. Biol. Chem. 259, 7747–7754.

25. Iwami, G., Kawabe, J., Ebina, T., Cannon, P.J., Homcy, C.J., and Ishikawa, Y. (1995). Regulation of adenylyl cyclase by protein kinase A. J. Biol. Chem. 270, 12481–12484. 10.1074/jbc.270.21.12481.

26. Hadcock, J.R., and Malbon, C.C. (1988). Down-regulation of β-adrenergic receptors: agonist-induced reduction in receptor mRNA levels. Proc. Natl. Acad. Sci. U. S. A. 85, 5021–5025. 10.1073/pnas.85.14.5021.

27. Collins, S., Bouvier, M., Bolanowski, M.A., Caron, M.G., and Lefkowitz, R.J. (1989). cAMP stimulates transcription of the β2-adrenergic receptor gene in response to short-term agonist exposure. Proc. Natl. Acad. Sci. U. S. A. 86, 4853–4857. 10.1073/pnas.86.13.4853.

28. Drube, J., Haider, R.S., Matthees, E.S.F., Reichel, M., Zeiner, J., Fritzwanker, S., Ziegler, C., Barz, S., Klement, L., Filor, J., et al. (2022). GPCR kinase knockout cells reveal the impact of individual GRKs on arrestin binding and GPCR regulation. Nat. Commun. 13, 540. 10.1038/s41467-022-28152-8.

29. Garbett, K.A., Zheng, C., Drube, J., Hoffmann, C., Gurevich, V.V., and Sando, R.C. (2026). Cytoplasmic tail diversity determines the effector bias of the adhesion GPCR ADGRL2. Cell Chem. Biol. 33, 364–378.e6. 10.1016/j.chembiol.2026.02.004.

30. Veldhuis, J.D., Iranmanesh, A., Mulligan, T., and Pincus, S.M. (1999). Disruption of the young-adult synchrony between luteinizing hormone release and oscillations in follicle-stimulating hormone, prolactin, and nocturnal penile tumescence (NPT) in healthy older men. J. Clin. Endocrinol. Metab. 84, 3498–3505. 10.1210/jcem.84.10.6100.

31. Upton, T.J., Zavala, E., Methlie, P., Kämpe, O., Tsagarakis, S., Øksnes, M., Bensing, S., Vassiliadi, D.A., Grytaas, M.A., Botusan, I.R., et al. (2023). High-resolution daily profiles of tissue adrenal steroids by portable automated collection. Sci. Transl. Med. 15, eadg8464. 10.1126/scitranslmed.adg8464.

32. Genter, P., Berman, N., Jacob, M., and Ipp, E. (1998). Counterregulatory hormones oscillate during steady-state hypoglycemia. Am. J. Physiol. 275, E821–829. 10.1152/ajpendo.1998.275.5.E821.

33. Schöfl, C., Becker, C., Prank, K., von zur Mühlen, A., and Brabant, G. (1997). Twenty-four-hour rhythms of plasma catecholamines and their relation to cardiovascular parameters in healthy young men. Eur. J. Endocrinol. 137, 675–683. 10.1530/eje.0.1370675.

34. Shannahoff-Khalsa, D.S., Kennedy, B., Yates, F.E., and Ziegler, M.G. (1996). Ultradian rhythms of autonomic, cardiovascular, and neuroendocrine systems are related in humans. Am. J. Physiol.-Regul. Integr. Comp. Physiol. 270, R873–R887. 10.1152/ajpregu.1996.270.4.R873.

35. Cannon, W.B. (1915). Bodily Changes in Pain, Hunger, Fear and Rage: An Account of Recent Researches Into the Function of Emotional Excitement (Appleton).

36. Ménard, L., Ferguson, S.S.G., Zhang, J., Lin, F.-T., Lefkowitz, R.J., Caron, M.G., and Barak, L.S. (1997). Synergistic Regulation of β2-Adrenergic Receptor Sequestration: Intracellular Complement of β-Adrenergic Receptor Kinase and β-Arrestin Determine Kinetics of Internalization. Mol. Pharmacol. 51, 800–808. 10.1124/mol.51.5.800.

37. McGraw, D.W., and Liggett, S.B. (1997). Heterogeneity in beta-adrenergic receptor kinase expression in the lung accounts for cell-specific desensitization of the beta2-adrenergic receptor. J. Biol. Chem. 272, 7338–7344. 10.1074/jbc.272.11.7338.

38. Violin, J.D., DiPilato, L.M., Yildirim, N., Elston, T.C., Zhang, J., and Lefkowitz, R.J. (2008). beta2-adrenergic receptor signaling and desensitization elucidated by quantitative modeling of real time cAMP dynamics. J. Biol. Chem. 283, 2949–2961. 10.1074/jbc.M707009200.

39. Purvis, J.E., Karhohs, K.W., Mock, C., Batchelor, E., Loewer, A., and Lahav, G. (2012). p53 dynamics control cell fate. Science 336, 1440–1444. 10.1126/science.1218351.

40. Albeck, J.G., Mills, G.B., and Brugge, J.S. (2013). Frequency-modulated pulses of ERK activity transmit quantitative proliferation signals. Mol. Cell 49, 249–261. 10.1016/j.molcel.2012.11.002.

41. Nelson, D.E., Ihekwaba, A.E.C., Elliott, M., Johnson, J.R., Gibney, C.A., Foreman, B.E., Nelson, C., See, V., Horton, C.A., Spiller, D.G., et al. (2004). Oscillations in NF-κB signaling control the dynamics of gene expression. Science 306, 704–708. 10.1126/science.1099962.

42. Hoffmann, A., Levchenko, A., Scott, M.L., and Baltimore, D. (2002). The IκB-NF-κB signaling module: temporal control and selective gene activation. Science 298, 1241–1245. 10.1126/SCIENCE.1071914.

43. Tsai, H.-C., Zhang, F., Adamantidis, A., Stuber, G.D., Bonci, A., de Lecea, L., and Deisseroth, K. (2009). Phasic firing in dopaminergic neurons is sufficient for behavioral conditioning. Science 324, 1080–1084. 10.1126/science.1168878.

44. Stavreva, D.A., Wiench, M., John, S., Conway-Campbell, B.L., McKenna, M.A., Pooley, J.R., Johnson, T.A., Voss, T.C., Lightman, S.L., and Hager, G.L. (2009). Ultradian hormone stimulation induces glucocorticoid receptor-mediated pulses of gene transcription. Nat. Cell Biol. 11, 1093–1102. 10.1038/ncb1922.

45. Wildt, L., Hausler, A., Marshall, G., Hutchison, J.S., Plant, T.M., Belchetz, P.E., and Knobil, E. (1981). Frequency and amplitude of gonadotropin-releasing hormone stimulation and gonadotropin secretion in the rhesus monkey. Endocrinology 109, 376–385. 10.1210/endo-109-2-376.

46. Oakley, R.H., Laporte, S.A., Holt, J.A., Barak, L.S., and Caron, M.G. (1999). Association of β-arrestin with G protein-coupled receptors during clathrin-mediated endocytosis dictates the profile of receptor resensitization. J. Biol. Chem. 274, 32248–32257. 10.1074/jbc.274.45.32248.

47. Goh, L.K., and Sorkin, A. (2013). Endocytosis of receptor tyrosine kinases. Cold Spring Harb. Perspect. Biol. 5, a017459. 10.1101/cshperspect.a017459.

48. Cohen-Saidon, C., Cohen, A.A., Sigal, A., Liron, Y., and Alon, U. (2009). Dynamics and Variability of ERK2 Response to EGF in Individual Living Cells. Mol. Cell 36, 885–893. 10.1016/j.molcel.2009.11.025.

49. Frick, C.L., Yarka, C., Nunns, H., and Goentoro, L. (2017). Sensing relative signal in the Tgf-β/Smad pathway. Proc. Natl. Acad. Sci. 114, E2975–E2982. 10.1073/pnas.1611428114.

50. Lee, R.E.C., Walker, S.R., Savery, K., Frank, D.A., and Gaudet, S. (2014). Fold change of nuclear NF-κB determines TNF-induced transcription in single cells. Mol. Cell 53, 867–879. 10.1016/j.molcel.2014.01.026.

51. Haney, S., and Hancox, R.J. (2005). Rapid onset of tolerance to β-agonist bronchodilation. Respir. Med. 99, 566–571. 10.1016/j.rmed.2004.10.014.

52. Salpeter, S.R., Ormiston, T.M., and Salpeter, E.E. (2004). Meta-analysis: respiratory tolerance to regular β2-agonist use in patients with asthma. Ann. Intern. Med. 140, 802–813. 10.7326/0003-4819-140-10-200405180-00010.

53. Wang, W.C.H., Mihlbachler, K.A., Brunnett, A.C., and Liggett, S.B. (2009). Targeted transgenesis reveals discrete attenuator functions of GRK and PKA in airway β2-adrenergic receptor physiologic signaling. Proc. Natl. Acad. Sci. U. S. A. 106, 15007–15012. 10.1073/pnas.0906034106.

54. Deshpande, D.A., Theriot, B.S., Penn, R.B., and Walker, J.K.L. (2008). β-Arrestins specifically constrain β2-adrenergic receptor signaling and function in airway smooth muscle. FASEB J. Off. Publ. Fed. Am. Soc. Exp. Biol. 22, 2134–2141. 10.1096/fj.07-102459.

55. Li, X., Burnight, E.R., Cooney, A.L., Malani, N., Brady, T., Sander, J.D., Staber, J., Wheelan, S.J., Joung, J.K., McCray, P.B., et al. (2013). piggyBac transposase tools for genome engineering. Proc. Natl. Acad. Sci. U. S. A. 110, E2279–E2287. 10.1073/pnas.1305987110.

56. Dobin, A., Davis, C.A., Schlesinger, F., Drenkow, J., Zaleski, C., Jha, S., Batut, P., Chaisson, M., and Gingeras, T.R. (2013). STAR: ultrafast universal RNA-seq aligner. Bioinformatics 29, 15–21. 10.1093/bioinformatics/bts635.

57. Liao, Y., Smyth, G.K., and Shi, W. (2014). featureCounts: an efficient general purpose program for assigning sequence reads to genomic features. Bioinformatics 30, 923–930. 10.1093/bioinformatics/btt656.

58. Lowe, D.G. (2004). Distinctive image features from scale-invariant keypoints. Int. J. Comput. Vis. 60, 91–110. 10.1023/B:VISI.0000029664.99615.94.

59. Stringer, C., and Pachitariu, M. (2025). Cellpose3: one-click image restoration for improved cellular segmentation. Nat. Methods 22, 592–599. 10.1038/s41592-025-02595-5.

60. Ershov, D., Phan, M.-S., Pylvänäinen, J.W., Rigaud, S.U., Le Blanc, L., Charles-Orszag, A., Conway, J.R.W., Laine, R.F., Roy, N.H., Bonazzi, D., et al. (2022). TrackMate 7: integrating state-of-the-art segmentation algorithms into tracking pipelines. Nat. Methods 19, 829–832. 10.1038/s41592-022-01507-1.

61. Rohatgi, A. WebPlotDigitizer, version 5.2. https://automeris.io/.

62. Feinstein, W.P., Zhu, B., Leavesley, S.J., Sayner, S.L., and Rich, T.C. (2012). Assessment of cellular mechanisms contributing to cAMP compartmentalization in pulmonary microvascular endothelial cells. Am. J. Physiol.-Cell Physiol. 302, C839–C852. 10.1152/ajpcell.00361.2011.

63. Conti, M., Mika, D., and Richter, W. (2014). Cyclic AMP compartments and signaling specificity: Role of cyclic nucleotide phosphodiesterases. J. Gen. Physiol. 143, 29–38. 10.1085/jgp.201311083.

64. Jockers, R., Angers, S., Da Silva, A., Benaroch, P., Strosberg, A.D., Bouvier, M., and Marullo, S. (1999). β2-adrenergic receptor down-regulation. Evidence for a pathway that does not require endocytosis. J. Biol. Chem. 274, 28900–28908. 10.1074/jbc.274.41.28900.

65. Seibold, A., January, B.G., Friedman, J., Hipkin, R.W., and Clark, R.B. (1998). Desensitization of β2-adrenergic receptors with mutations of the proposed G protein-coupled receptor kinase phosphorylation sites. J. Biol. Chem. 273, 7637–7642. 10.1074/jbc.273.13.7637.

66. Cullum, S.A., Veprintsev, D.B., and Hill, S.J. (2023). Kinetic analysis of endogenous β2-adrenoceptor-mediated cAMP GloSensor^TM^ responses in HEK293 cells. Br. J. Pharmacol. 180, 1304–1315. 10.1111/bph.16008.

67. Freedman, N.J., Liggett, S.B., Drachman, D.E., Pei, G., Caron, M.G., and Lefkowitz, R.J. (1995). Phosphorylation and desensitization of the human β1-adrenergic receptor: Involvement of G protein-coupled receptor kinases and cAMP-dependent protein kinase. J. Biol. Chem. 270, 17953–17961. 10.1074/jbc.270.30.17953.

68. Clark, R.B., and Perkins, J.P. (1971). Regulation of adenosine 3’:5’-cyclic monophosphate concentration in cultured human astrocytoma cells by catecholamines and histamine. Proc. Natl. Acad. Sci. U. S. A. 68, 2757–2760. 10.1073/pnas.68.11.2757.

69. Marsh, J.D., and Roberts, D.J. (1987). Adenylate cyclase regulation in intact cultured myocardial cells. Am. J. Physiol. 252, C47–54. 10.1152/ajpcell.1987.252.1.C47.

70. Sibley, D.R., Peters, J.R., Nambi, P., Caron, M.G., and Lefkowitz, R.J. (1984). Desensitization of turkey erythrocyte adenylate cyclase. β-adrenergic receptor phosphorylation is correlated with attenuation of adenylate cyclase activity. J. Biol. Chem. 259, 9742–9749. 10.1016/S0021-9258(17)42762-1.

71. Frangakis, M.V., and Kimelberg, H.K. (1985). Desensitization of β-receptors on primary astrocyte cultures by norepinephrine but not by tricyclic antidepressants. Brain Res. 339, 49–56. 10.1016/0006-8993(85)90620-1.

72. Refsnes, M., Sandnes, D., and Christoffersen, T. (1987). The relationship between beta-adrenoceptor regulation and beta-adrenergic responsiveness in hepatocytes. Studies on acquisition, desensitization and resensitization of isoproterenol-sensitive adenylate cyclase in primary culture. Eur. J. Biochem. 163, 457–466. 10.1111/j.1432-1033.1987.tb10891.x.

73. Galant, S.P., and Britt, S. (1984). Uncoupling of the beta-adrenergic receptor as a mechanism of in vitro neutrophil desensitization. J. Lab. Clin. Med. 103, 322–332. 10.5555/uri:pii:0022214384901793.

74. Tougui, Z., Do Khac, L., and Harbon, S. (1980). Modulation of cyclic AMP content of the rat myometrium: desensitization to isoproterenol, PGE2 and prostacyclin. Mol. Cell. Endocrinol. 20, 17–34. 10.1016/0303-7207(80)90091-x.

75. Yu, S.S., Lefkowitz, R.J., and Hausdorff, W.P. (1993). β-adrenergic receptor sequestration. A potential mechanism of receptor resensitization. J. Biol. Chem. 268, 337–341. 10.1016/S0021-9258(18)54155-7.

76. Trovero, F., Marin, P., Tassin, J.P., Premont, J., and Glowinski, J. (1994). Accelerated resensitization of the D1 dopamine receptor-mediated response in cultured cortical and striatal neurons from the rat: respective role of alpha 1-adrenergic and N-methyl-D-aspartate receptors. J. Neurosci. 14, 6280–6288. 10.1523/JNEUROSCI.14-10-06280.1994.

